# Simulating Iron Deficiency in Plant Plastidial Metabolism With a Flexible Neural-Mechanistic Hybrid Approach

**DOI:** 10.1101/2025.06.10.658179

**Authors:** Sara El Alaoui, Christopher S. Henry, Crysten Blaby, Tim Paape, Meng Xie, Samuel M. D. Seaver

## Abstract

Flux balance analysis has proven to be a successful approach in metabolic engineering and systems biology to predict intracellular fluxes of large genome-scale networks and the essentiality of genes encoding enzymes and regulatory factors. Flux balance analysis (FBA) relies on a key assumption of a metabolic state being persistent (“steady”) over a given time frame. This assumption works well for microbial growth because of the ease with which microbial media can be fixed, biomass can be decomposed, and growth rates can be measured. However, the assumption is far less tenable for the cells and tissues of complex multicellular organisms, particularly if any integrated data is sampled from a heterogeneous collection of developing cells continually interacting between and across tissues. These will likely exhibit transient metabolic states equilibrating over varying timescales, and many FBA studies in complex organisms typically either ignore time as a parameter, or integrate data taken over long timescales (days/weeks). In this work, we adopt and modify a previously published machine learning approach originally developed to hybridize several aspects of a constraint-based approach with machine-learning in order to predict growth. This approach accommodates transient state dynamics, at the cost of tolerating a controlled amount of slack in the steady-state assumption, in order to enable transcript-constrained flux estimation in plant tissues without optimizing a growth objective. For our case study, we reconstruct the metabolism of the plastid of Poplar and Sorghum, integrating data sampled from leaf tissue under varying levels of iron bioavailability. The two species diverge sharply: Sorghum suppresses its photosynthetic electron transport and Calvin cycle abruptly at day 7 and its carbon delivery to biomass collapses to a third of control by day 21, whereas Poplar declines gradually and retains roughly 70%. Normalizing each reaction score to the plastid proteome pool makes the two species comparable and exposes reallocation as the pool contracts, including a split within Sorghum’s sulfate assimilation that tracks which enzymes carry iron cofactors.

## 1. INTRODUCTION

The state of a metabolic network is the set of constituent metabolites and their abundances at any given point in time. Understanding how metabolic states are relevant to the fitness and function of an organism requires us to establish how the metabolic state may persist or change in a given environment (Grimbs et al., 2007; Ratcliffe & Shachar-Hill, 2006; Rontein et al., 2002; Sajitz-Hermstein & Nikoloski, 2016; Stephanopoulos & Vallino, 1991). Flux Balance Analysis is one of several computational approaches that has been developed to explore the range of metabolic states that are possible within a large metabolic network representing the cumulative work of hundreds of enzymes (Orth et al., 2010). The success of the approach is contingent on the key assumption of establishing a reasonable time frame for any predicted metabolic state as steady-state.

As such, the approach was first developed for the exponential growth phase of microbes where the growth rate itself can be assumed to be invariant until stationary phase is reached (Varma et al., 1993; Varma & Palsson, 1993a, 1993b).

In order for the FBA approach to reliably predict a metabolic state that is valid given the experimental conditions, researchers need to be able to define the external constraints of the metabolic network, namely its inputs and outputs, and the time frame within which the experimental data was gathered. This typically includes the media as the inputs, and the biomass and excreted metabolites as the outputs. Given the ease with which this data can be captured for any microbe, as well as its growth rate, it follows that FBA has been successful for generating reliable predictions under a number of environmental conditions for different microbial species (Feist & Palsson, 2008; Gianchandani et al., 2010; O’Brien et al., 2015; Price et al., 2003). The success of using FBA (and other approaches reliant on stoichiometric matrices) as a predictive approach means that high-throughput phenotyping using arrays such as those produced by Biolog (Microbial Phenotype Assay, 2023) can be used to explore how changes in the media (inputs) can affect the output (growth/biomass) (Feist et al., 2007; Henry et al., 2017; Oberhardt et al., 2008; Oh et al., 2007; Zingue et al., 2017). The resulting data can either be integrated back into metabolic reconstructions to improve the predictions, used as validation, or, in the case of Faure et al. and Tazza et al., used to train an ML model (Faure et al., 2023; Tazza et al., 2025).

Stoichiometric approaches have also been applied to the genome-scale metabolic reconstructions for complex multicellular organisms, notably humans (Swainston et al., 2016; Robinson et al., 2020) and plants (Cheung et al., 2013; Cunha et al. 2023; Sampaio et al., 2024). Researchers focus on narrowing the focus of their modeling approach by integrating data from specific tissues and conditions, such as the liver in humans (Mardinoglu et al., 2014), or the leaf in plants (Cheung et al., 2014). Even in cases where researchers attempt to model multicellular organisms at a larger scale, they still modularize their approach around individual tissues (Gomes de Oliveira Dal’Molin et al., 2015; Martins Conde et al., 2021). The approaches taken when tackling multicellular organisms have been successful where researchers compare their results by integrating data from different experimental conditions, but for the same tissue, reconstructed network, and timepoint. For example, in the case of the human liver, Mardinoglu et al. analyze results generated from the liver of different patients to identify potential metabolic markers for a liver disease. In the case of the plant leaf, Cheung et al. compare results generated from a leaf during the day/night cycle to identify differential fluxes during C3 and CAM photosynthesis.

The global mass-balance constraint of stoichiometric approaches ensures that there is a direct mechanistic link between the metabolites chosen for the inputs (media) and outputs (biomass) of the simulations, and the activity of the internal metabolic network (Orth et al., 2010). For this reason, researchers will try to establish as complete a set of metabolites that comprise the boundary, through the literature or through their own sampling of the metabolites in the experiments. However complete the set of metabolites are, the fact remains that the pathways that are activated during a simulation are contingent on the formation of the media and the biomass. Consequently, a given simulation may utilize only a fraction of the reconstructed metabolic network. This is particularly problematic when integrating omics data that would indicate that some enzymes and pathways would be active, but the underlying reactions are blocked because key metabolites are not incorporated in the biomass (Yuan et al., 2016).

In recent years, there has been a small but growing number of ML approaches that have been used with FBA to solve a number of biological problems in a few microbial species (Choudhury et al., 2024; Isewon et al., 2024; Kugler & Stensjö, 2024; Song et al., 2025; Vĳayakumar et al., 2020; Wu et al., 2024; Wendering & Nikoloski, 2026). In each of these cases, the steady-state solutions generated by running FBA were used as inputs to train the machine-learning approach, and in turn were dependent on the composition of the media and the biomass. In two recent studies that build on each other, the researchers develop an approach that allowed the flux solution to be reached using an optimization approach other than flux balance analysis (Faure et al., 2023; Tazza et al., 2025). Faure et al. applied AMN-QP, a gradient descent approach for optimizing the metabolic fluxes that met the mass-balance constraint but is not as restrictive as applying FBA with linear programming where a blocked reaction can shut down the entire network. Building on this work, Tazza et al. (2025) proposed MINN, a metabolic-informed neural network that extends AMN-QP to integrate multi-omics data. MINN first uses a data-driven feed-forward neural network to map transcriptomics, proteomics, and two measured exchange fluxes corresponding to glucose and oxygen to an initial GEM-wide flux estimate, *V*_0_. Then, similar to AMN-QP, *V*_0_ is refined through multiple gradient descent iterations minimizing a mechanistic FBA-based loss. The full MINN model is trained with a custom loss that combines the data-driven error between predicted and measured fluxes with FBA constraint violations.

While MINN has shown superior performance compared to pFBA and simple ML methods, its mechanistic layer is more complex than it needs to be for our purpose: the constraints can be applied directly through FBA penalty terms in a single loss function. Our departure from MINN is therefore architectural rather than a claim to better supervision — we use no measured fluxes at all. In our approach, transcript abundance sets a flux ceiling for each reaction, and gradient descent moves the fluxes only as far as mass balance requires without optimizing growth. Here we explore the extension of the approach by Faure et al. to integrate transcriptomics data with the steady-state constraints of FBA offering a simpler and more straightforward formulation, while still bringing together the same elements they aim to combine through bi-level optimization. By applying the steady-state constraints of FBA as “soft” constraints, this allows the approach to fit fluxes through active enzymes and pathways in a manner that is uncoupled across the network to varying degrees. This will lead to predictions that are not completely mass-balanced but the areas of imbalance can follow enzymatic activity and can be informative. The approach allows us to explore beyond the fraction of the metabolic network responsible for the biosynthesis of the biomass components and discover metabolic activity in other pathways.

We should be clear at the outset about what is and is not neural about this. We inherit the AMN architecture and its physics-informed loss, but in the present study we operate it in *solver* mode: no weights are trained, and the procedure reduces to gradient descent on a constrained optimization problem, initialized at the transcript-derived flux vector. We adopt this formulation not because a neural network is required to solve the underlying problem — it is not — but because casting the mechanistic constraints as a differentiable loss keeps the door open to using that same loss as a trainable layer once datasets large enough to train on become available, which is the direction the AMN family was built for.

In order to test our approach we focus on the smaller but mostly complete metabolic reconstruction of the plant plastid for several reasons. The biosynthetic network of the plant plastid is (almost) solely responsible for a large number of key primary biomass components (Rolland et al., 2018), and, due to its directly acquired source of carbon assimilated via Rubisco, and energy assimilated by the photosynthetic electron transport chain, we are able to reduce the solution space and take a closer look at how our results indicate what the plastid needs to consume and produce in order to achieve its function. It is also interesting because the plastid (as well as the mitochondrion) is microbial in origin, and has independent growth rates, in that it doubles at a rate separate from that of the parent cell, although there is evidence that the plant tissues have mechanisms for increasing/slowing plastidial growth rates (Miyagishima, 2011).

We also test our approach by comparing datasets generated from identical experiments for two different plant species. Following the trend of using the curated reconstruction of plant generalized metabolism for different flowering plant species as part of the PlantSEED project (Seaver et al., 2014, 2018), we are able to project the same metabolic reconstruction of the plant plastid onto different plant species for which data has been sampled under identical conditions. The Quantitative Plant Science Initiative (QPSI) at Brookhaven National Laboratory (Xie et al., 2024) carried out experiments where they grew Poplar and Sorghum seedlings in hydroponics with normal and low levels of iron over several weeks (Mishra et al., 2025). The seedlings exhibited the classical signs of chlorosis, and transcript abundances were sampled from the leaves over time. By applying our transient-state modeling approach to these datasets, we reconstruct the dynamic metabolic shifts occurring within the plastids of both species. Our results reveal a systemic reprioritization of the plastidial network, highlighting specific iron-binding and ferredoxin-dependent pathways that dynamically uncouple from standard biomass expectations as the plant shifts resources from general growth to photosynthesis.

## 2. METHODS

We summarize our approach in Figure 1 and as follows. We start with the reconstruction of plant generalized metabolism for Arabidopsis thaliana as generated from the data made available by the PlantSEED project (Seaver et al., 2014, 2018). The curation of the Arabidopsis enzymes for the project has been extended to include phylloquinone biosynthesis and NADP+-malate dehydrogenase since its last publication. From this reconstruction, we extract the plastidial network. We modify the plastidial metabolic reconstruction to ensure it can produce flux and biosynthesize many primary biomass components. We use OrthoFinder protein families to assign the Poplar and Sorghum genes encoding the same enzymes as the curated Arabidopsis genes. We develop a novel integrative process for estimating enzyme abundances from transcriptomics, using the RNASeq dataset generated by the QPS Initiative. Finally, we outline our hybrid ML-FBA approach, and present results on the simulations in the plastidial metabolic reconstructions.

**Figure 1.**
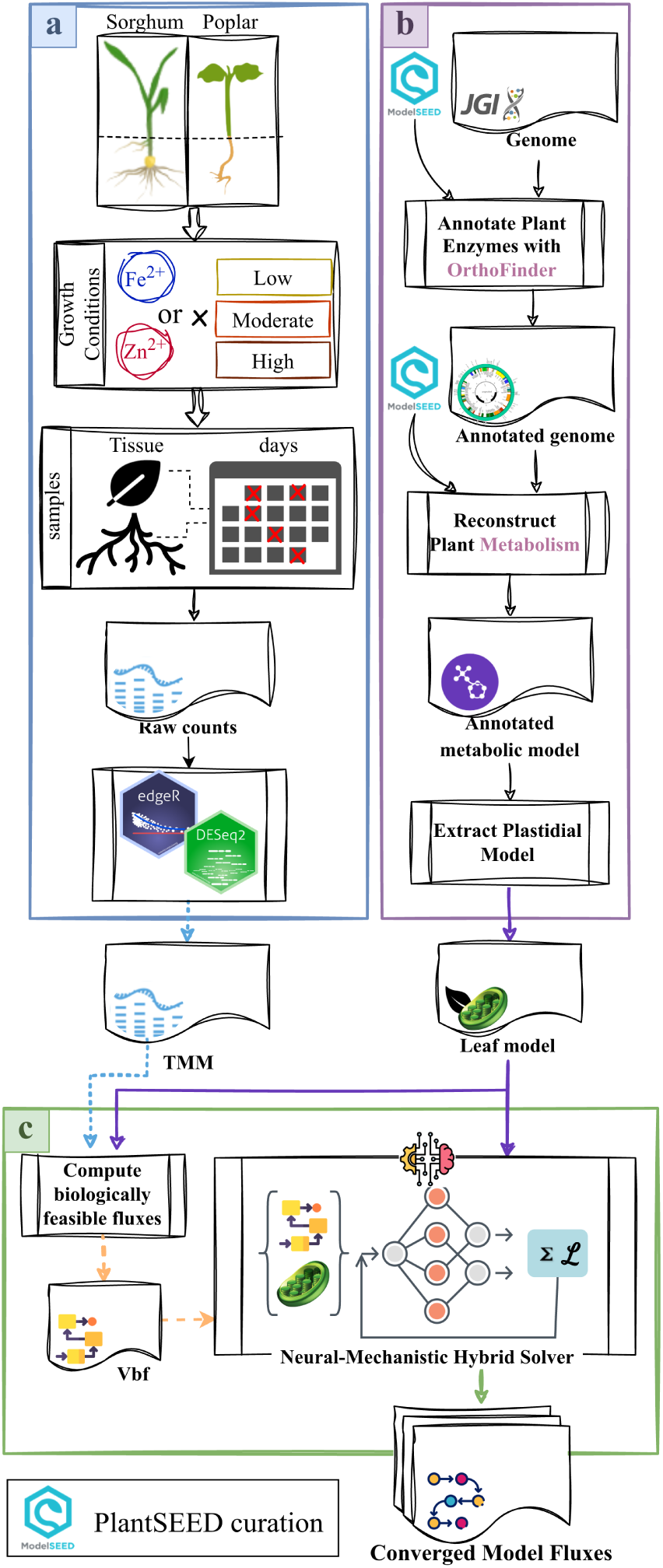
General overview of our approach. Our work was generally carried out in three steps: **(a)** The parent QPSI experiment grew triplicates of P. trichocarpa (Torr. & Gray) Nisqually-1 and S. bicolor cv. BTx623 under low, moderate and high availability of two metals (iron and zinc). Leaf and root tissues of the control triplicates were sampled at 0hr, with an additional sampling of all treatments’ root tissue at 1hr. Henceforth, all replicates were sampled at 2 days, 4 days, 7 days, 14 days and 21 days. RNA sequencing of the samples was performed using Illumina short-read technology. Raw counts were then processed to compute Trimmed Mean of M-values (TMM) and log2 fold change. TMM-normalized abundances were averaged across the three biological replicates before scoring, so each condition contributes a single abundance per transcript. **The present study uses a subset of this design**: leaf tissue only, moderate iron (Control, 10 µM Fe) versus low iron (FeLim, 0 µM Fe), at the five day-scale timepoints plus the 0hr Control baseline — 11 conditions per species (§3.3). The processed dataset is then integrated with the metabolic reconstruction of the two species for the *in silico* simulations. **(b)** Enzymatic annotation and metabolic reconstruction for each species. We use the reference genomes for Sorghum (v3.1.1) (McCormick et al., 2018) and Poplar (v4.1) (Tuskan et al., 2006). Here we generate predictions of enzymes encoded in the genomes using OrthoFinder (Emms & Kelly, 2019) v2.5.5, assigning the curated metabolic functions to protein-coding genes in Sorghum and Poplar. We also use the results to predict the protein-encoding genes that would localize to the plastid as part of our work on the reaction score and biologically feasible flux. To reconstruct the metabolic models, we utilize the mappings between the annotations and biochemical reactions that were curated for Arabidopsis thaliana, as part of the PlantSEED project (Seaver et al., 2014, 2018). As we limit the scope of this work to the plastidial network, we extract plastidial reactions from both species’ metabolic reconstructions. **(c)** In this last step, we simulate the plastidial metabolic reconstructions with the TMM dataset using the neural-mechanistic hybrid model, AMN. We start by computing the biologically feasible flux (*V_b__f_*), which is used to constrain the simulated fluxes for each condition. The AMN-derived solver uses FBA constraints together with the *V_b__f_*values to obtain a converged model flux vector for each condition and treatment.

### 2.1. Species

The two species used in this work, *Populus trichocarpa* (Poplar) (Tuskan et al., 2006) and *Sorghum bicolor* (Sorghum) (McCormick et al., 2018), are bioenergy crops whose genomes have been championed by Joint Genome Institute (Nordberg et al., 2014). Poplar is a perennial flowering tree native to North America in the family Salicaceae. It is considered to have the most mature reference assembly for perennial plants of interest as a bioenergy crop (Tuskan et al., 2006). Sorghum is an annual C4 grass in the family Poaceae that is grown in the Southeast and Midwest of the USA as a bioenergy crop. Sorghum is known to be tolerant to drought and soil degradation (Brenton et al., 2016; McCormick et al., 2018). The protein FASTA files for both genomes were downloaded from Phytozome (Goodstein et al., 2011). We used reference genome v4.1 for Poplar and v3.1.1 for Sorghum.

Our work was developed during a collaborative effort with the Quantitative Plant Science Initiative (Xie et al., 2024), and here we use a dataset for both Poplar and Sorghum individuals grown under control and limited-iron conditions (Mishra et al., 2025). Triplicate plants of Poplar and Sorghum were hydroponically grown under two conditions: Control with moderate iron bioavailability (10 µM Fe) and low Fe (FeLim) at 0 µM. For Poplar, leaves with the same leaf plastochron index (LPI = 4) were sampled. For Sorghum, the second youngest leaf was sampled. Leaf tissues were sampled for RNA sequencing at 2 days, 4 days, 7 days, 14 days and 21 days. RNASeq data was processed for the three replicates in the two growth conditions. The final step in processing the transcript abundances was to use the Trimmed Mean of M-values (TMM) normalization approach (Robinson & Oshlack, 2010) to normalize the abundances of the transcripts across the samples for each species. The TMM-normalized abundances of the three biological replicates were then averaged within each species/tissue/treatment/timepoint combination, so that every condition contributes a single transcript-abundance vector to the reaction-score calculation (§2.4) and, in turn, a single simulated condition.

### 2.2. Modeling Plastidial Metabolism

We use the full reconstruction of plant generalized metabolism available online as part of the PlantSEED project. For the curation of the protein localization data, the PlantSEED curators used several resources such as PPDB (Sun et al., 2009) and SUBA (Hooper et al., 2022) to determine if the proteins would localize to the plastid. From this curation, we can extract the plastidial sub-network of the Arabidopsis metabolic reconstruction. We compose a plastid-specific media providing some necessary metabolites that the plastid is not able to biosynthesize on its own. We build a plastid-specific biomass reaction, consisting of the primary biomass components that the plastid is able to biosynthesize and export for the growth and maintenance of the plant tissue. Finally, we fill several gaps in the metabolic reconstruction for some pathways that are not entirely contained within the plastid. After the plastidial network was finalized, we used Flux Variability Analysis (FVA) to find and remove blocked transport reactions. To ensure consistency, we compared FBA and FVA results obtained using both the KBase platform and the COBRApy package. Even though the plastid incorporates only nearly 28% of the reactions found in the full reconstruction of plant generalized metabolism, it can biosynthesize 52% of the biomass components found in the full metabolic reconstruction. The working plastidial reconstruction is available in our repository (see Section 5).

### 2.3. Annotation of Enzymes

We use OrthoFinder (Emms & Kelly, 2019) v2.5.5 to generate plant protein families from the protein FASTA files of 20 different species as downloaded from Phytozome (Goodstein et al., 2011) and to predict orthologs of the Arabidopsis genes. For each enzyme-encoding gene in Arabidopsis, we project its enzymatic function onto a gene in Sorghum and Poplar if they follow one of the two conditions: (1) They must be predicted orthologs with at least 30% pairwise sequence identity and (2) they must otherwise be in the same protein family with a pairwise sequence identity greater than the average of the orthologs from the same family. Of the reactions in the plastidial metabolic reconstruction for which an Arabidopsis gene was curated to encode an enzymatic function, we find that 95.1% are predicted to be associated with Sorghum genes and 97.7% are predicted to be associated with Poplar genes. The two plastidial metabolic reconstructions for Poplar and Sorghum are found in our repository (see Section 5).

### 2.4. Reaction Scores

In order to convey how the TMM-normalized abundances of individual transcripts could affect the overall flux of the reaction, we follow a similar logic as previously published (Seaver et al., 2015) for identifying the subunit that limits the formation of the enzymatic complex albeit with a difference. In the new formulation, if there are a number of paralogs for a single subunit, we compute the sum of abundances of all the paralogs instead of taking the abundance of the paralog with the highest expression. This sum is then used as the abundance of the subunit in question, before comparing it to other subunits. We summarize our approach in Figure 2.

**Figure 2.**
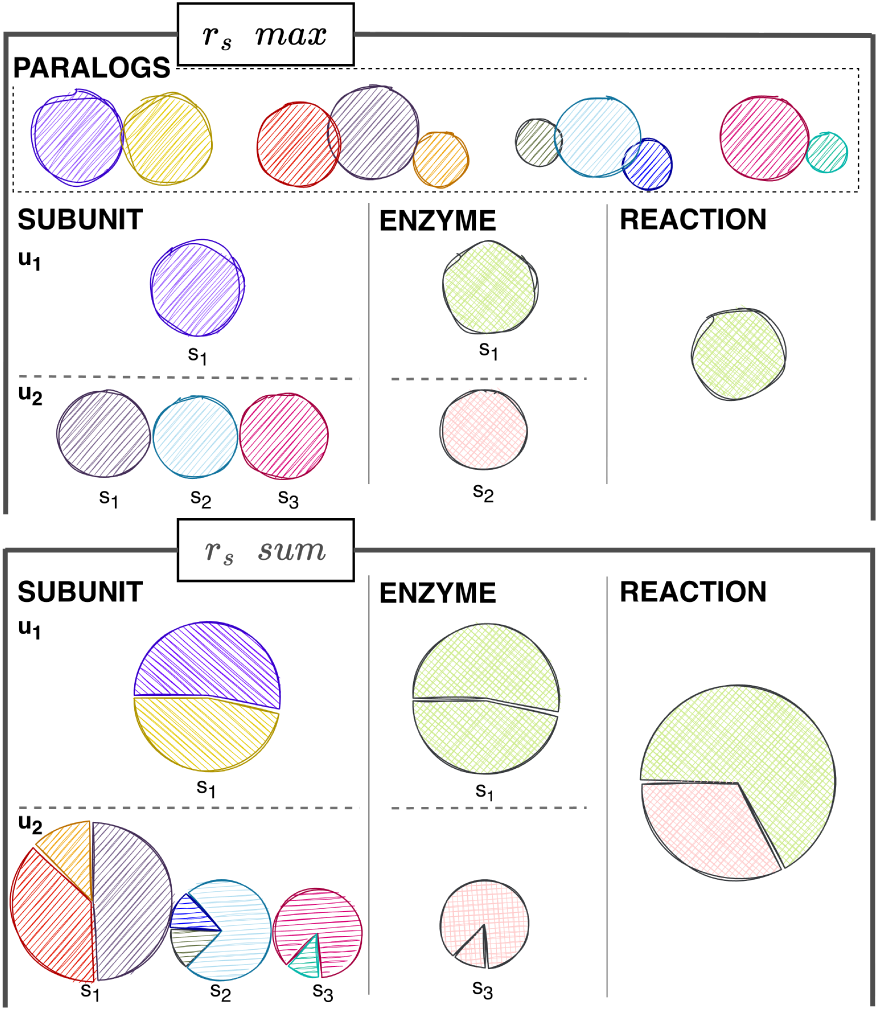
Calculating reaction score as an estimate of enzyme abundance. A graphic visualization of how the abundances of transcripts that encode different subunits may contribute to the overall estimated abundance of enzymatic complexes, and in turn, how the abundance of one or more enzyme may contribute to our estimation of the upper limit of a reaction’s flux (see main text), which we call the reaction score. In this example, we address the Acetyl-CoA carboxyl transferase activity for which two distinct and multi-subunit enzymes are encoded in the Arabidopsis genome (Lamesch et al., 2011). The data used to generate the figure were taken from the leaf of a Poplar grown after three weeks on limited iron. Furthermore, each subunit may have multiple paralogs, especially given the different ploidy in the genomes of different plant species. The TMM-normalized abundance of every transcript is shown as part of a pie-chart for each step in our logic in determining the reaction score, and we also show three different approaches. At the top we show the old approach from Seaver et al. (2015), *r_s_max*, in the middle we show the updated approach where the abundances of paralogs that encode the same subunit are added cumulatively, *r_s_sum*, and in the bottom we show the same approach but normalized by the size of the proteome in the plastid, *r̃_s_sum*,(see main text). We show that the different approaches may result in a different upper bound for the flux of the enzymatic activity in the metabolic reconstruction.

As illustrated in Figure 2, the score of reaction *i* in Eq. 1 is the sum of the complex expression scores (*u _j_*) of all enzymatic complexes (*U_j_*) that catalyze it. As any of the enzymatic complexes can catalyze a reaction, we sum their complex expression scores rather than take the maximum as in Seaver et al. (2015). The complex expression score of an enzymatic complex, *u _j_*, is the minimum score of its constituent subunit scores, *s_k_* (Equation 2). Because an enzymatic complex requires all constituent subunits to be present, its activity is limited by the least abundant subunit. Finally, the subunit score, *s_k_*, is computed using Eq. 3 as the sum of the TMM-normalized transcript abundance (*p_l_*) for all paralogs in the set that carries out the function of the subunit *p_k_*. While we find that plant generalized metabolism is highly conserved and there are very few cases where an ortholog for a nuclear-encoded protein subunit was not predicted, we also understand that a number of important and highly-expressed protein complexes involved in photosynthesis, the Calvin cycle, and respiration, contain subunits encoded in the two respective organellar genomes, and we do not expect to find these subunits encoded in the nuclear genomes of other plant species. To this end, we maintain a short list of these subunits by name, and we exclude them from the computation of the reaction score. We understand that there’s an underlying assumption of correlation between transcript and protein abundances, and this is a limitation of this method.

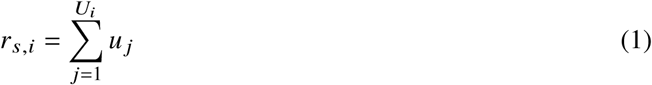

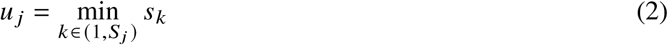

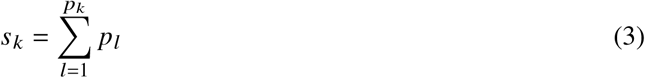

The two species differ substantially in the size of their plastid proteome pools. We define the total plastid proteome *P_total_* as the summed transcript abundance of all enzymes whose Arabidopsis orthologs are localized to the plastid — collected from two localization sources (Bouchnak et al., 2019; Sun et al., 2009) and mapped to Poplar and Sorghum orthologs (*p_plastid_*) with OrthoFinder — computed per species, timepoint, and growth condition (Eq. 4).

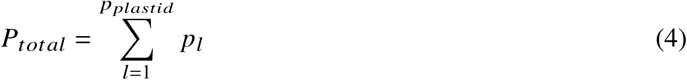

Table 1 reports the mean plastid- versus non-plastid transcript abundance per species and treatment: Poplar carries more matched plastidial orthologs than Sorghum (2,489 vs. 2,008), yet Sorghum’s plastid genes are expressed at roughly twice the level, so the plastid proteome pool *P_total_* differs ∼1.7× between species and contracts under iron limitation in both. Figure 3 shows the plastid- versus non-plastid abundance distributions over the time course and the number of reactions whose objective- versus relative-score response reaches the 95th-percentile significance threshold, which peaks at day 7 for Sorghum and day 21 for Poplar.

**Figure 3.**
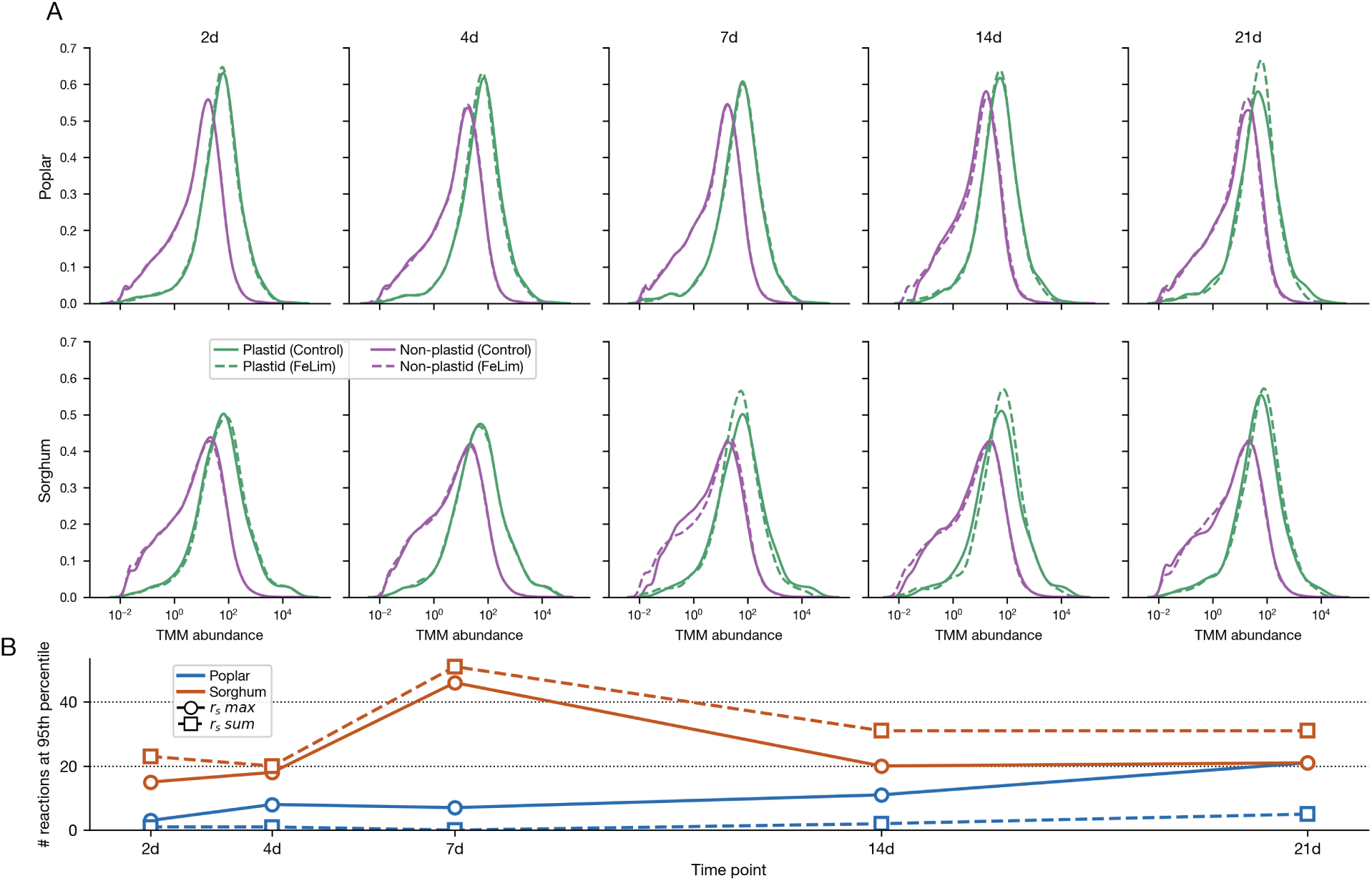
Abundance of the plastid proteome is affected by FeLim from day 7 onward in Sorghum and at day 21 in Poplar. Panel A shows the probability distributions of transcript abundance for proteins localized in the plastid against non-plastidial transcripts, demonstrating that the higher peaks at day 7 (Sorghum) and day 21 (Poplar) are specific to the iron limitation response in the plastid. In panel B we show the number of reactions that fall in the 95th percentile from Figure 5, highlighting the different temporal responses between the two species in plastidial metabolism: the Sorghum count peaks at day 7 and remains elevated through days 14 and 21, whereas the Poplar count rises late and peaks at day 21.

**Table 1.** Mean TMM-normalized expression of plastid-localized vs. non-plastid genes by species and treatment. Poplar has more matched plastidial orthologs than Sorghum, Sorghum plastidial transcripts have nearly twice the abundance.

| Treatment | Species | Plastid mean | Non-plastid mean | Fold change |
| --- | --- | --- | --- | --- |
| Control | Poplar | 212.65 | 27.27 | 7.80× |
|  | Sorghum | 467.93 | 31.94 | 14.65× |
|  | *Fold change* | 2.20× | 1.17× | — |
| FeLim | Poplar | 185.10 | 25.71 | 7.20× |
|  | Sorghum | 368.18 | 31.13 | 11.83× |
|  | *Fold change* | 1.99× | 1.21× | — |

To compare enzyme investment *across* species despite this pool-size difference, we express each reaction score relative to the plastid proteome pool, defining the plastid-relative reaction score *r̃_s_*_,*i*_ by normalizing *r_s_*_,*i*_ by *P_total_* (Eq. 5).

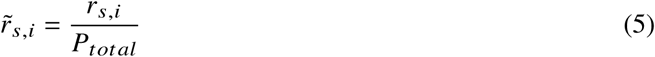

We use the plastid-relative reaction score *only* for the cross-species comparison of metabolic allocation (section 3.4.3). **Caution:** the relative score must not be used to generate or interpret flux. Dividing by the contracting plastid pool decouples the score from enzyme abundance and inflates iron-limited targets, so fluxes produced from *r̃_s_* cannot be read quantitatively. Accordingly, the apparent catalytic rate (*K_app_*, §2.5.3) and the biologically feasible flux (*V_b f_*, §2.6) are computed from the objective reaction score *r_s_*, not *r̃_s_*.

### 2.5. Apparent Catalytic Rate

The apparent catalytic rate (*K_app_*) is the estimation of the catalytic rate of enzymes *in vivo* (Davidi et al., 2016; Valgepea et al., 2013). Davidi et al. (2016) describe how they use data from a number of growth experiments (30) to estimate the highest possible catalytic rate for each enzyme in *Escherichia coli*. We consider the plastid to be embedded within a relatively consistent environment (compared to microbes) in the cytosol of plant cells. We do not have growth rates for the plastid, so we take a different approach to the estimation of the highest apparent catalytic rate. We employed flux variability analysis (FVA) alongside a novel constraint-based method called Flexible Biomass. By allowing individual biomass components to vary relative to one another within appropriate physiological bounds, this approach maximizes the feasible solution space and captures the widest possible range of fluxes. This builds on earlier analyses of the metabolic plasticity underlying biomass-precursor production (Shaw & Kundu, 2015), and reflects a concept similar to the ensemble biomass representations of Choi et al. (2023).

#### 2.5.1. Splitting Reversible Reactions

Model reactions, including media exchange reactions, need to be irreversible from left to right in order for all fluxes to be positive (Eq. 15). As such, any reversible reaction in the model is converted into two opposing reactions both irreversible from left to right. For each pair of split forward and reverse reactions with fluxes *v _f_* and *v_r_*, respectively, we load the FVA-derived maximum flux for each reaction and compute the net maximum flux across the pair. If the net flux is nonzero, the pair is resolved into a single effective direction using eqs. (6) and (7), which sets the reaction with the lower flux to 0 and deducts the lower flux from the greater flux.

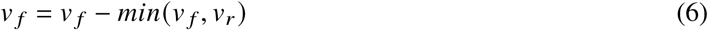

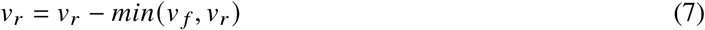

If the net flux is 0 and the reaction score is above 0, the pair is treated as a *futile cycle* — a split reaction whose forward and reverse copies both carry flux, circulating substrate with no net conversion (an artifact of the splitting of reversible reactions, distinct from an ATP-driven biochemical futile cycle). In this case, both directions are retained, but assigned a representative median flux so that the reactions are not artificially blocked. This median is calculated from the model-wide flux distribution after excluding reactions with artificially high futile-cycle fluxes. When a median flux is assigned to both directions of a split pair, this value is split equally across the two reactions to preserve the total capacity constraint. Lastly, if a reaction is blocked in the FVA analysis and has a reaction score greater than 0, it is assigned the median flux to avoid artificially blocking it.

A pair of split forward and reverse reactions can both saturate at high futile-cycle fluxes, and their net value may then be close to zero simply because two solver-ceiling values differ by numerical noise rather than by a real capacity difference. To distinguish this case from a genuine directional signal — for example, a Calvin-cycle reaction where one direction has above-ceiling capacity and the other does not — we require both criteria to be satisfied before flagging a pair. The original pFVA maxima for both directions (see §2.5.3) must be at least 90% of the futile-cycle ceiling, and the two values must be within 1% of each other in magnitude. Pairs meeting both tests are treated as ceiling-saturated: we reset both directions to zero and let the both-zero-plus-scored rule described in the previous paragraph lead to them being assigned the median flux. Without this step, arbitrary differences would lead to a *V_b f_* being assigned to one direction and not the other. The correction reclassified 137 split pairs in Poplar and 134 in Sorghum.

#### 2.5.2. Flexible Biomass

With FBA, or any approach that utilizes a stoichiometric matrix coupled with vectors for media and for biomass, the coefficient for each biomass component is fixed according to the experimental data of the species in question. Given the uncertainty surrounding not only the biomass of the plastid itself but the exported metabolites from the plastid that comprise the biomass of the surrounding plant cell and tissue, we did not want to restrict our approach to a set of fixed coefficients for biomass components. While some approaches typically allow for some degree of flexibility in the coefficients, we go a step further and permit substantially greater flexibility, allowing biomass coefficients to be neutralized (coefficient set to, or close to zero) or even to be reversed, indicating that the biomass component could be consumed rather than exported. We have developed a set of additional variables and constraints that may be added to a metabolic model to support a more flexible formulation of the composition of the biomass objective function. In the most basic version of this formulation, we first introduce a bi-directional drain reaction for every metabolite already included in the biomass objective function, or that is being considered for inclusion in it. Each such drain adds one new flux variable, *V_bio_*___*_comp_*_,*i*_, carrying flux through the bi-directional reaction *bio*_*comp*, *i* ⇌ θ, whose sign convention is given below.

Next, we add constraints setting bounds on the flux through these drain reactions *V_bio_*___*_comp_*_,*i*_, Eq. 8:

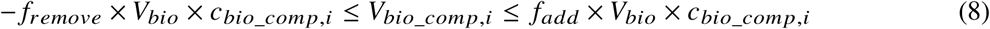

Where *f_remove_* is a fraction between 0 and 1 indicating the fraction of *bio*_*comp*, *i* that can be removed from the biomass reaction; *f_add_* is a fraction between 0 and 1 indicating the fraction of *bio*_*comp*, *i* that can be added to the biomass reaction; and *c_bio_*___*_comp_*_,*i*_ is the stoichiometric coefficient of *bio*_*comp*, *i* in the biomass reaction. Note that the bounds on the biomass component drain fluxes cannot be fixed, because they must vary with the flux through the biomass reaction itself. Nevertheless, these remain simple linear constraints. Lastly, we add constraints that ensure that the total mass of the biomass reaction is preserved despite changes in biomass composition caused by pushing flux through the biomass component drain fluxes, Eq. 9.

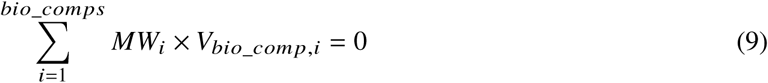

where *MW_i_* is the molecular weight of biomass component *i*. Generally, one will want to minimize the sum of *V_bio_*___*_comp_*_,*i*_ to ensure that only the minimal needed changes are made to the biomass reaction. Once a solution is obtained, negative *V_bio_*___*_comp_*_,*i*_ values indicate a decrease in the biomass composition of component *i*, while positive values indicate an increase.

The flexible biomass package is implemented as part of the suite of CobraPy-compatible functions that have been developed in ModelSEEDPy (https://github.com/ModelSEED/ModelSEEDPy) and can be applied to any metabolic model loaded as a Cobra model.

#### 2.5.3. K_app_ Computation

The hybrid ML-FBA approach uses integrated FBA-like stoichiometric constraints to explore the activation of pathways within the space of biologically feasible fluxes; therefore, we use parsimonious Flux Variability Analysis (pFVA; Lewis et al., 2010) with the ModelSEEDpy flexible biomass package to generate and explore as broad a solution of interdependent fluxes as possible. We run pFVA with a fraction-of-optimum of 0.75 (requiring the objective to remain at ≥ 75% of its FBA optimum) and a pFBA factor of 1.2 (allowing each reaction’s maximum to explore up to 1.2× the parsimonious minimum). Even with the pFBA penalty, reactions embedded in futile cycles still saturate at a solver-imposed ceiling; the downstream correction we apply to those reactions is described in §2.5.1. We find the maximal flux (*MF_i_*) that reaction *i* can achieve and use this value to estimate the highest apparent catalytic rate for the reaction, *K_app_*_,*i*_, in equation 10.

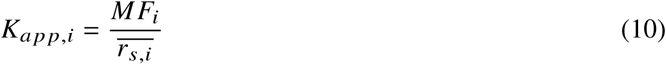

Where *r_s_*_,*i*_ is the average of the objective reaction score derived from the data collected for the control experiments for reaction *i* over all the time points.

This mirrors the construction of Davidi et al. (2016), whose Eq. 2 likewise divides a flux-balance solution by an enzyme abundance; the reaction score stands in for their proteomic copy numbers as our estimate of enzyme abundance.

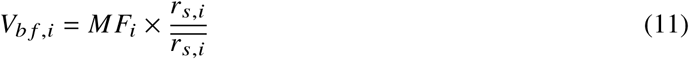

that is, each reaction’s capacity ceiling is its pFVA maximum scaled by the fold-change of its reaction score relative to the control mean. A reaction whose transcripts are at control levels is capped at its pFVA maximum; one whose transcripts have halved is capped at half of it.

### 2.6. Biologically Feasible Flux

In this work, we build on the approach of Faure et al. by adopting and modifying their hybrid ML-FBA framework. A key distinction of our approach is the incorporation of transcriptomics data, which was not included in their original framework. We use transcriptomics data to estimate the range of the flux for each reaction in the network that is considered to be feasible *in vivo*.

The biologically feasible flux (*V_b f_*) of a reaction *i* during any experimental condition and at any time point in the dataset is computed using Equation 12.

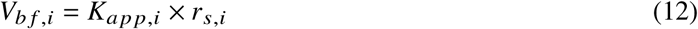

As explained in section 2.5.1, some metabolic reactions are assigned an *MF_i_* value of zero. If their objective reaction score *r_s_*_,*i*_ > 0 (indicating the enzyme is active), we set *MF_i_* to a representative median calculated from the model-wide FVA flux distribution after excluding reactions with artificially high futile-cycle fluxes. This assigns a nonzero *V_b f_* _,*i*_ to the reaction, even if it is blocked when running FBA, allowing the optimizer to activate it, subject to the activity of its catalyzing enzyme. Finally, the biologically feasible flux is the value that we use to constrain fluxes for each reaction when we apply our Physics-informed Neural Network approach.

### 2.7. Neural-Mechanistic Hybrid Model

We adopt a hybridization of constraint-based modeling and ML called the Artificial Metabolic Network (AMN) (Faure et al., 2023). AMN is an application of the Physics-informed Neural Network (PiNN) approach to metabolic modeling. PiNNs are founded on the training of ML models using problem-specific differential equations within custom loss functions. In doing so, PiNNs overcome the limitations of small datasets and guide predictions using domain knowledge. Hence, in this implementation, the model follows the constraints of FBA to improve model predictions in lieu of fitting data. Such models can operate as predictors given a learning dataset or as solvers when data is unavailable or limited. We depict the workflow of applying the integrated data as part of our modeling approach and generating a set of converged fluxes in Figure 4.

**Figure 4.**
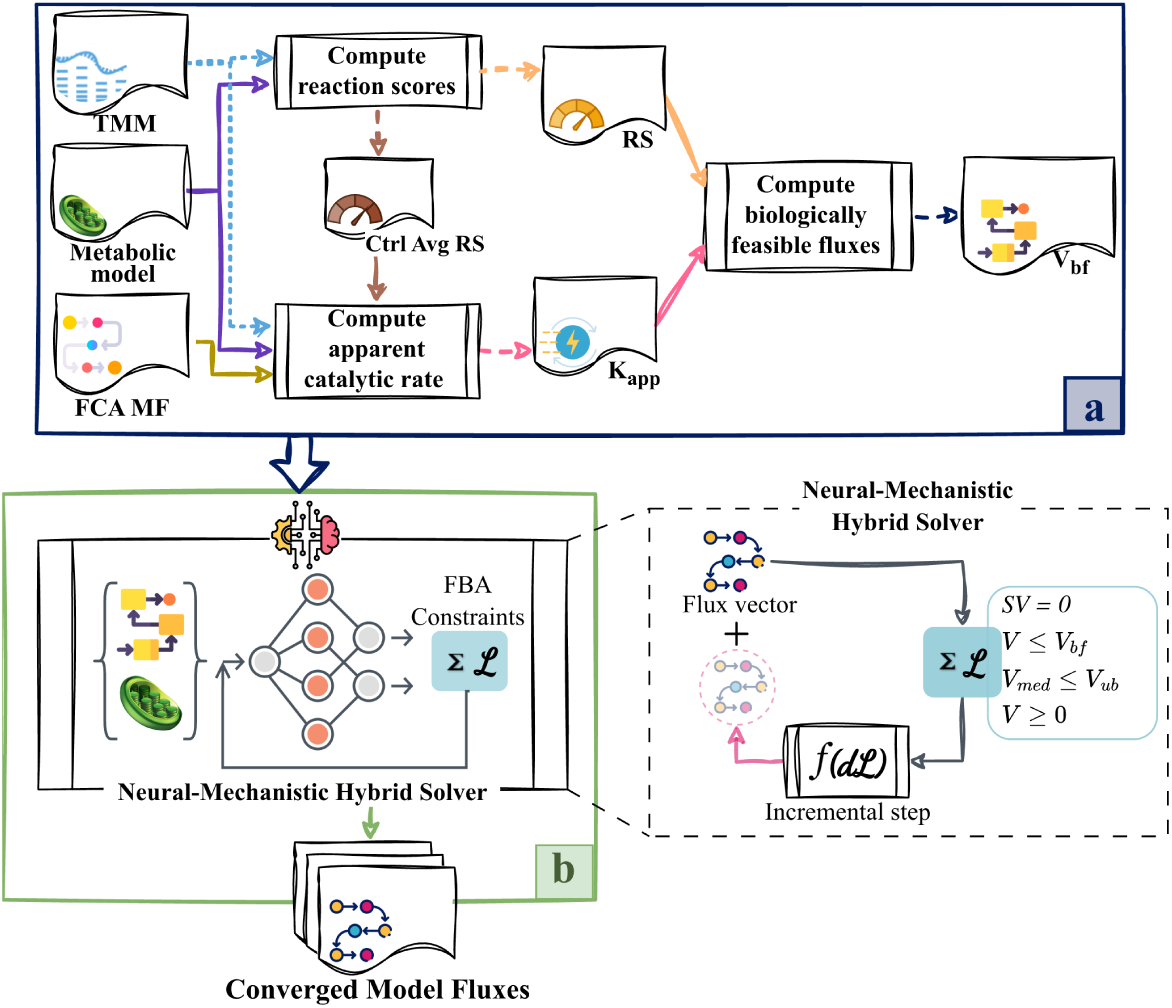
Workflow of the neural-mechanistic hybrid approach. Graphic visualization of the workflow for the approach developed as part of the present work. a) From the integration of TMM-normalized abundances to the computation of the control-averaged objective reaction score for estimation of the apparent catalytic rate, and the per-condition objective reaction score for the estimation of the upper bound of the biologically feasible flux for each reaction. b) The biologically feasible flux is integrated with a gradient descent solver that mirrors the metabolic reconstruction of the plastid, along with a set of constraints representing the Flux Balance Analysis approach that define the loss function. The approach iteratively generates a set of fluxes and incrementally steps down the loss gradient for several thousand iterations until a solution is reached.

AMN is designed to function as a prediction model that trains on a relatively small dataset of media components and bacterial growth rates; obtaining such datasets for plants is difficult for several reasons, including and not limited to, different tissues and organs developing at different rates, shifts between developmental and growth stages, and varying environmental cues. However, AMN allows custom loss functions and can be used as a QP/LP solver when training data is not available. In this work, we modify and extend AMN in three key aspects outlined in the subsequent sections: i) integration of biologically feasible flux; ii) customized loss functions; and iii) projection and bound enforcement.

#### 2.7.1. Integration of V_b f_

The hybrid AMN approach combines FBA constraints, the stoichiometry of the mechanistic model, and the predictive capabilities of machine learning. Similarly to hybrid AMN (Faure et al., 2023), our model incorporates several constraints borrowed from FBA (Equations 13–15): the fundamental steady state constraint, a constraint on positive flux and constraints on media uptake. Additionally, we include a new constraint that sets an upper limit on the reaction fluxes based on experimental transcriptomic data (Eqs. 16 and 17). As detailed in the formulation and explained in Section 2.6, we compute each reaction’s *V_b f_* using the approximate abundance of the enzyme catalyzing the reaction. We then use the *V_b f_* value as an upper limit for the flux that could be carried by a specific reaction under each treatment and at each time point. The split forward and reverse reactions are limited by the same underlying enzyme capacity so their combined simulated flux is constrained as the sum of the two directions and evaluated against the assigned bidirectional capacity, *V_b f_* . Besides serving as this upper bound, *V_b f_* is also the initial condition, with internal reactions initialized at *V* ^(0)^ = *V_b f_* and media exchanges at their *V_med_* values. Since every loss term is one-sided (§2.7.2) and is therefore minimized by *V* = **0**, the converged fluxes are best read as a projection of the transcript-derived vector onto the set of mass-balanced flux distributions rather than as the optimum of a biological objective.

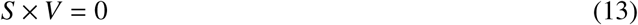

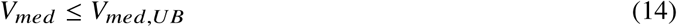

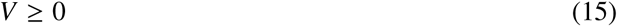

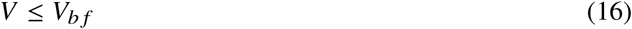

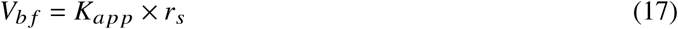

#### 2.7.2. Customized loss function and early stopping

Briefly described, a loss function is used by ML models to evaluate the accuracy of their predictions. A loss function, which typically computes the difference between predicted and expected values, guides the learning process with the aim of minimizing this difference. A custom loss function is designed with the same overarching goal, but it is problem specific and may not necessarily rely on standard functions like Root Mean Squared Error (RMSE) or others. The custom loss function used in our method is derived from the formulation in eqs. (13) to (17). By design, this function does not rely on expected values (i.e. ground truth) to measure accuracy. Instead, it focuses on satisfying the transcriptome-based stoichiometric constraints. As a result, the hybrid mechanistic ML method proposed here serves as a solver rather than a predictor.

The loss function returns a total loss, *Loss_t_*, defined in Eq. 23. *Loss_t_* incurred by the model is calculated as the sum of five individual losses, each defined as follows. Media upper bounds and positive flux constraints are defined in Equations 18 and 19, respectively. *P_med_* is the *n_med_* × *n* projection matrix defined in Faure et al. (2023) where each matrix element corresponding to a media exchange reaction is set to 1. Equation 20 is the steady state constraint loss with the addition of a penalty, *p*, to better enforce it. We define the loss function corresponding to the biologically feasible flux constraint in Equation 21. The nominal transcriptderived capacity *V_b f_* defines the target upper bound used in the loss function (Eq. 21). Because this bound is enforced softly through the loss, individual gradient-descent updates may transiently exceed *V_b f_* . To prevent larger excursions, after each update we apply the buffered clamp described in §2.7.3, which imposes an absolute ceiling of 1.10*V_b f_* . Finally, Equation 22 is a complementarity penalty — so called because it is zero exactly when at most one of each pair of complementary (forward/reverse) copies carries flux — that suppresses the *futile cycles* introduced by splitting reversible reactions (§2.5.1).

Throughout these expressions *n* is the number of reactions in the split network, *m* the number of metabolites, and *n_med_* the number of media exchange reactions; each squared residual is normalized by the number of rows it contains, i.e. 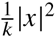 for a residual vector *x* of length *k*. *V_lb_* in eq. (19) is the vector of reaction lower bounds, which is zero for every reaction once reversible reactions have been split, so in practice 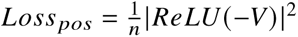.

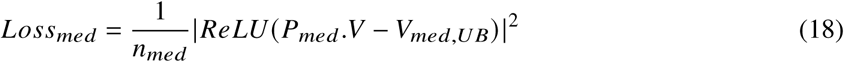

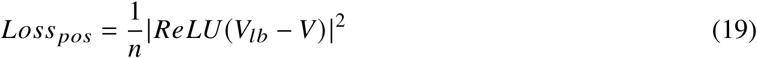

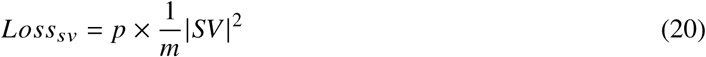

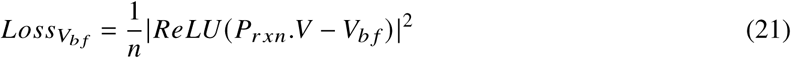

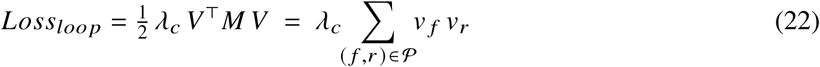

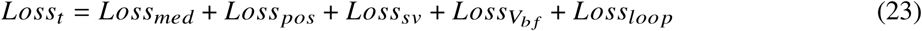

During the metabolic model annotation process, not all reactions are assigned an enzyme (see Section 2.3), which means that these reactions will not have an associated *V_b f_* . Therefore, we define *P_r xn_*, in the Eq. 21, as an *n* × *n* projection matrix where each element *X_ii_* is set to 1 if the corresponding reaction *r_i_* has a *v_b f_* value, and 0 otherwise. This matrix is used to indicate the presence of *v_b f_* values for each reaction in the system, thereby restricting *v_b f_* loss computation to this subset of reactions. *V_b f_* is the vector of size *n* where each element *o_i_* is set to the *V_b f_* value for reaction *i* if one is available, and 0 otherwise.

Because the hybrid mechanistic solver requires every flux to be non-negative, each reversible reaction is split into two one-way reactions — a forward copy *f* and a reverse copy *r* (§2.5.1). This split introduces an artifact: nothing prevents the optimizer from running *both* copies at once, converting substrate to product and immediately back again. Such a forward–reverse pair produces no net conversion and consumes nothing, yet it lets the optimizer satisfy the flux constraints “for free” by circulating arbitrarily large flux around the pair — the *futile cycle* defined in §2.5.1, and the source of the inflated fluxes noted there. Equation 22 describes a penalty that shuts down these futile cycles. The loss is the *product* of the fluxes for the two split reactions, *v _f_ v_r_*, summed over the set P of split forward/reverse pairs; *M* is the symmetric adjacency matrix that pairs them (*M _f r_* = *M_r f_* = 1, zero elsewhere). As a flux is never negative, this product is positive only when a reaction is split *and* both of its copies carry flux at the same time, so the penalty acts on exactly the futile cycles and nothing else: reactions that were never split, and split reactions running in a single net direction, are not penalized. Minimizing *v _f_ v_r_* (analytic gradient *λ_c_ MV*, added to the total-loss gradient at each step) drives one copy of every offending pair to zero, collapsing the cycle to a single, thermodynamically sensible net direction. We set the weight *λ_c_* = 0.01 — large enough to eliminate the futile-cycle flux, small enough to leave the mass-balance and *V_b f_* fits intact.

To further improve performance, we added an early stopping mechanism that monitors the loss behavior of each experimental condition independently during optimization. A relative-improvement threshold of 10^−2^ is applied per iteration to each condition’s loss term: if the loss does not drop by more than this fraction of its current best value, that condition’s patience counter is incremented; otherwise it is reset to zero. When a single condition’s patience counter exceeds the patience window (500 iterations), that condition is frozen by masking its row of the gradient matrix to zero, so its flux vector **V***_c_* stops updating while the remaining, still-improving conditions continue updating under the shared gradient-descent loop. The solver run as a whole terminates only once every condition has been frozen this way. This per-condition freezing prevents already-converged conditions from accumulating unnecessary updates – and from being dragged by gradients driven by other conditions still in transit – while preserving capacity for slower conditions to reach the same stable region.

#### 2.7.3. Projection and Bound Enforcement

Another key difference from Faure et al. (2023) is that our model does not maximize biomass and does not aim to fit predictions to any experimental data on uptake or individual biomass components. Unlike their framework, our method relaxes the transcript-derived flux vector onto the constraint set without forcing biomass growth. Instead of relying purely on the soft *V_b f_* loss term (Eq. 21) to enforce the per-reaction upper bound, we apply an additional proportional buffered clamp at the end of each gradient-descent step. For every reversible reaction pair whose combined capacity *V_f_* + *V_r_* exceeds 1.10 × *V_b f_*, both component fluxes are scaled back proportionally so that the pair sits at exactly 1.10 × *V_b f_* . The 10% buffer accommodates the residual uncertainty in the transcript-to-flux mapping while still preventing runaway growth on any single reaction. The clamp remains active throughout optimization, ensuring that no flux exceeds the permitted 10% buffer above *V_b f_* . In the converged flux vector, the largest excess reaches exactly this limit, indicating that at least one reaction pair saturates the ceiling while none exceeds it. The non-negativity projection (*V* ≥ 0) is applied unchanged. In so doing, “growth” and compound production in the model are regulated mainly by the steady-state and transcriptome-derived *V_b f_* constraints, which enables the hybrid approach to explore pathways that would otherwise be inactive. Lastly, this setup effectively avoids the added complexity of the multi-step loss computation introduced by Tazza et al. (2025), allowing the simulation to benefit from both data-driven and mechanistic constraints within single-stage iterations. Additional loss terms can also be incorporated as fluxomics or metabolomics data become available.

## 3. RESULTS

We present our results in four parts. We first report the orthologous-enzyme prediction across Arabidopsis, Sorghum, and Poplar (section 3.1) and the enzyme-level reaction scores and their response to iron limitation (section 3.2), then the statistical behavior of the gradient-descent optimizer (section 3.3), and finally the converged fluxes themselves (section 3.4).

### 3.1. Prediction of Orthologous Enzymes

We use OrthoFinder (Emms & Kelly, 2019) to predict the set of protein families that have proteins encoded by genes in Arabidopsis, Sorghum, and Poplar. We find that not all the orthologous proteins as predicted by OrthoFinder have a high sequence identity with the originally curated Arabidopsis enzyme, so we adopt a conservative approach for finding and predicting orthologous enzymes for Sorghum and Poplar in the metabolic reconstruction generated by this work. We find that our approach was able to assign enzymes to almost every reaction for which an Arabidopsis enzyme was curated (97.7% in Poplar and 95.1% in Sorghum).

### 3.2. Reaction Scores

We modify a previously published approach for estimating the abundance of the enzymes that are actively catalyzing flux in our simulations, as described in the methods. In Figure 5, we show how the reaction scores vary between the control and limited-iron datasets for every time point in the experimental dataset and for both species. The plastidial metabolic network is identical for the two species, and the sets of reactions for which we were able to compute a reaction score are near-identical — 95.1% of the enzyme-associated reactions are scored in Sorghum and 97.7% in Poplar (section 3.1) — allowing us to compare the results directly.

**Figure 5.**
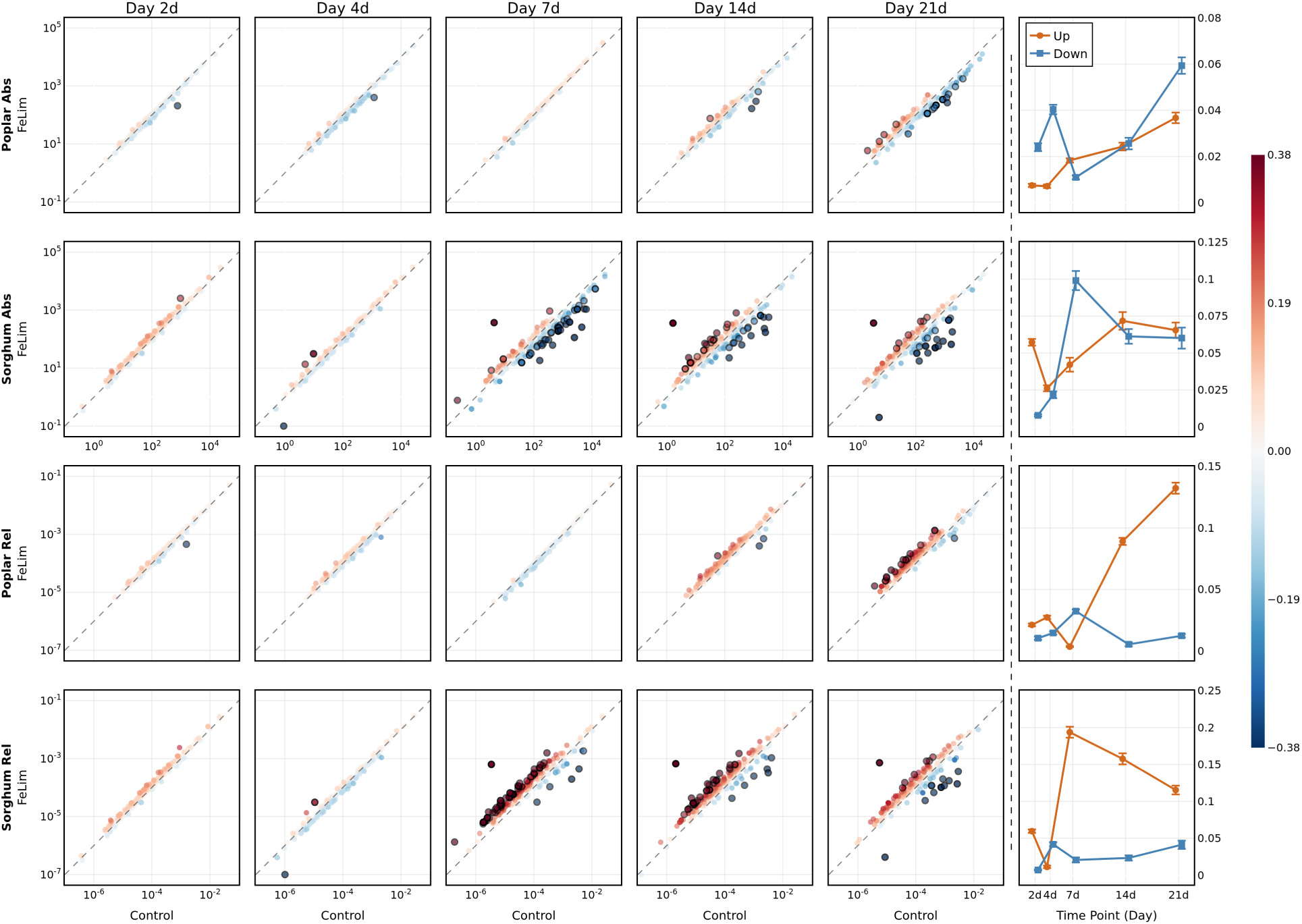
Objective and relative reaction scores of plastidial enzymes in Poplar and Sorghum in response to iron limitation. Each scatter plots the reaction score of an enzyme under FeLim vs Control conditions. Both axes are logarithmic, as reaction scores span roughly five orders of magnitude; the dashed diagonal is the identity line and I-dist is the perpendicular distance to it in log space, so it is a scaled log fold change and is independent of how large the two scores are. The first two rows are the objective reaction score *r_s_*, the last two rows are the relative reaction score *r̃_s_*. The color encodes I-dist (blue = down, red = up), and the top 5% by I-dist are outlined in black (see Figure 3). Column 6 gives the weighted mean and standard error of I-dist. Objectively, in rows 1-2, plastidial enzymes in Sorghum are displaced sharply downward at day 7, with a substantial subset remaining displaced through days 14 and 21, whereas in Poplar the response is muted until day 21. Relatively, in rows 3-4, both plastid proteomes drop in total abundance, leaving the enzymes of plastidial metabolism as the most abundant enzymes by day 7 in Sorghum and day 21 in Poplar.

For each reaction in the model for which there are TMM-normalized values assigned, instead of calculating the fold-change of *r_s_* between two conditions, we instead calculate the perpendicular distance (Ballantine & Jerbert, 1952) of the pair of reaction scores to the identity line, which we name I-dist. Because reaction scores are approximately log-normal across five orders of magnitude, we take this distance in log space, which makes I-dist a scaled log fold change: it indicates the change in the enzyme’s abundance between two conditions independently of the original size of the reaction scores. Both species exhibit more variation in the I-dist computed between the control and limited-iron conditions over time, and more so in Sorghum. The objective reaction score *r_s_* is what drives the transcript-constrained flux analyzed in the following sections (Figures 7 and 8), whereas the relative reaction score *r̃_s_* underpins the cross-species allocation comparison (Figure 9).

**Figure 6.**
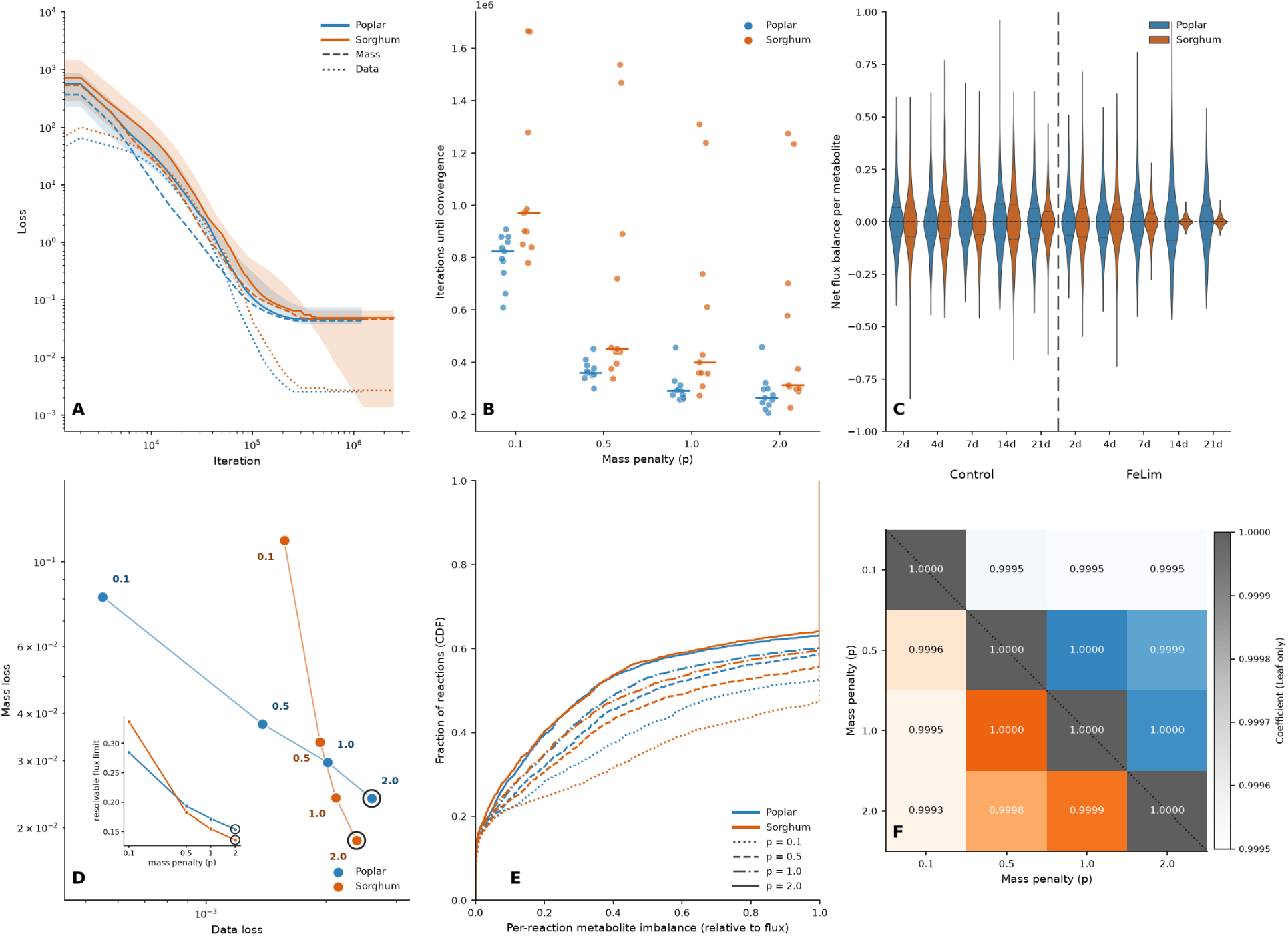
Dynamics of gradient-descent approach. Optimization convergence, early-stop dynamics, and stoichiometric violation diagnostics computed on data sampled from leaves under control and limited iron conditions. A) Evolution of the two key loss terms during gradient descent at the adopted operating point, mass penalty *p* = 2. The two per-term averages converge in a similar manner; the shaded band spans the 5th to 90th percentile of the total loss across conditions. B) Distribution of the number of steps per condition until early stopping is triggered, shown separately per mass penalty *p*. C) Per-condition imbalance violins at the operating point, showing how tightly the stoichiometric residual is compressed. D) The trade-off between *L*_data_ and *L*_mass_ across *p* identifies an elbow at *p* = 2.0 for both species (ringed marker), where the mass-balance residual is compressed at only a slight data-fit cost relative to *p* = 1.0. We therefore adopt *p* = 2.0 for both species; as panel F confirms, the bulk flux vectors at *p* = 1 and *p* = 2 remain highly correlated, so this choice sharpens mass balance without materially changing the reaction-level flux pattern. E) Cumulative distribution of |*SV* | per reaction across mass penalties; at *p* = 2.0 (solid) the CDF sits well below any biologically meaningful mass-balance threshold. F) Triangle-split heatmap of the Pearson correlation between flux vectors obtained at different mass-penalty values. Every off-diagonal correlation exceeds 0.98, and *p* = 1 vs. *p* = 2 correlates at *r* = 0.9996 (Sorghum) and *r* = 0.9997 (Poplar).

**Figure 7.**
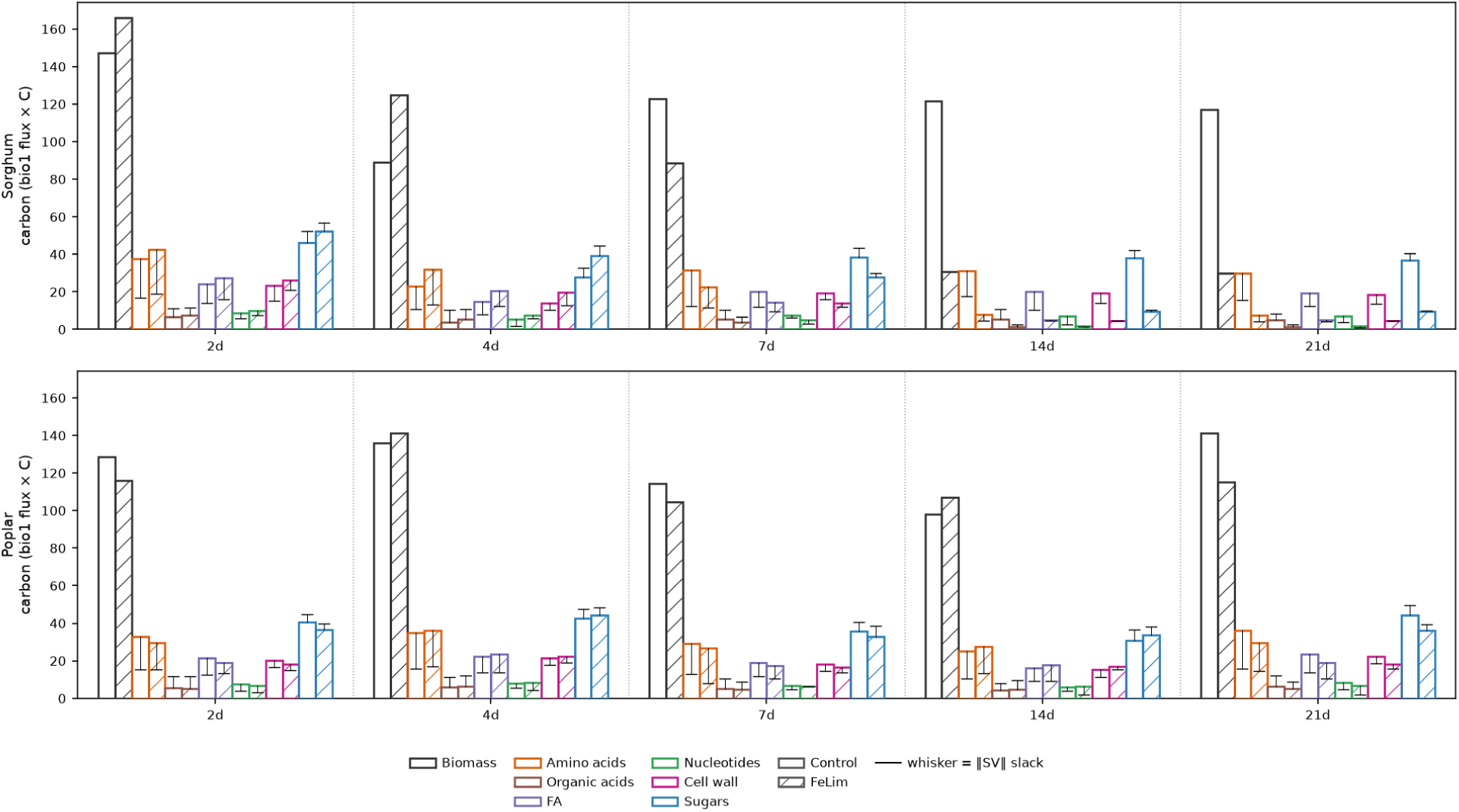
Transcript-driven carbon flux dynamically utilizes the biomass reaction as a drain. Each bar is the carbon drawn into the biomass reaction for one component class at one Leaf timepoint, in carbon atoms (bio1 flux × per-metabolite carbon number, summed over the class); the leftmost *Biomass* bar is the total over all classes. Solid outlines are Control, diagonal hatching is iron limitation (FeLim); rows are the two species, dotted lines separate timepoints. *Overall biomass:* total biomass carbon is high under Control and declines under FeLim, progressively over the time course and most severely by 14–21d, with a much larger drop in Sorghum than Poplar. The biomass reaction is not optimized; this is an emergent property indicating how much carbon the transcript-constrained plastidial network delivers (§3.4.1). Therefore, it falls when iron limitation curtails carbon fixation, and falls most in Sorghum. *Component classes:* the fixed biomass coefficients mean all classes scale together—sugars and amino acids dominate. *Whiskers* mark the carbon actually *supplied* to each class by the rest of the network (net production summed over all non-bio1 reactions). A whisker higher than the bar means the class is over-supplied and the excess is absorbed as mass-balance slack; a whisker *below* the bar means it is under-supplied, with the shortfall met by slack rather than pathway flux. Amino acids are consistently under-supplied (biosynthesis lags demand), whereas sugars and organic acids are consistently over-supplied.

**Figure 8.**
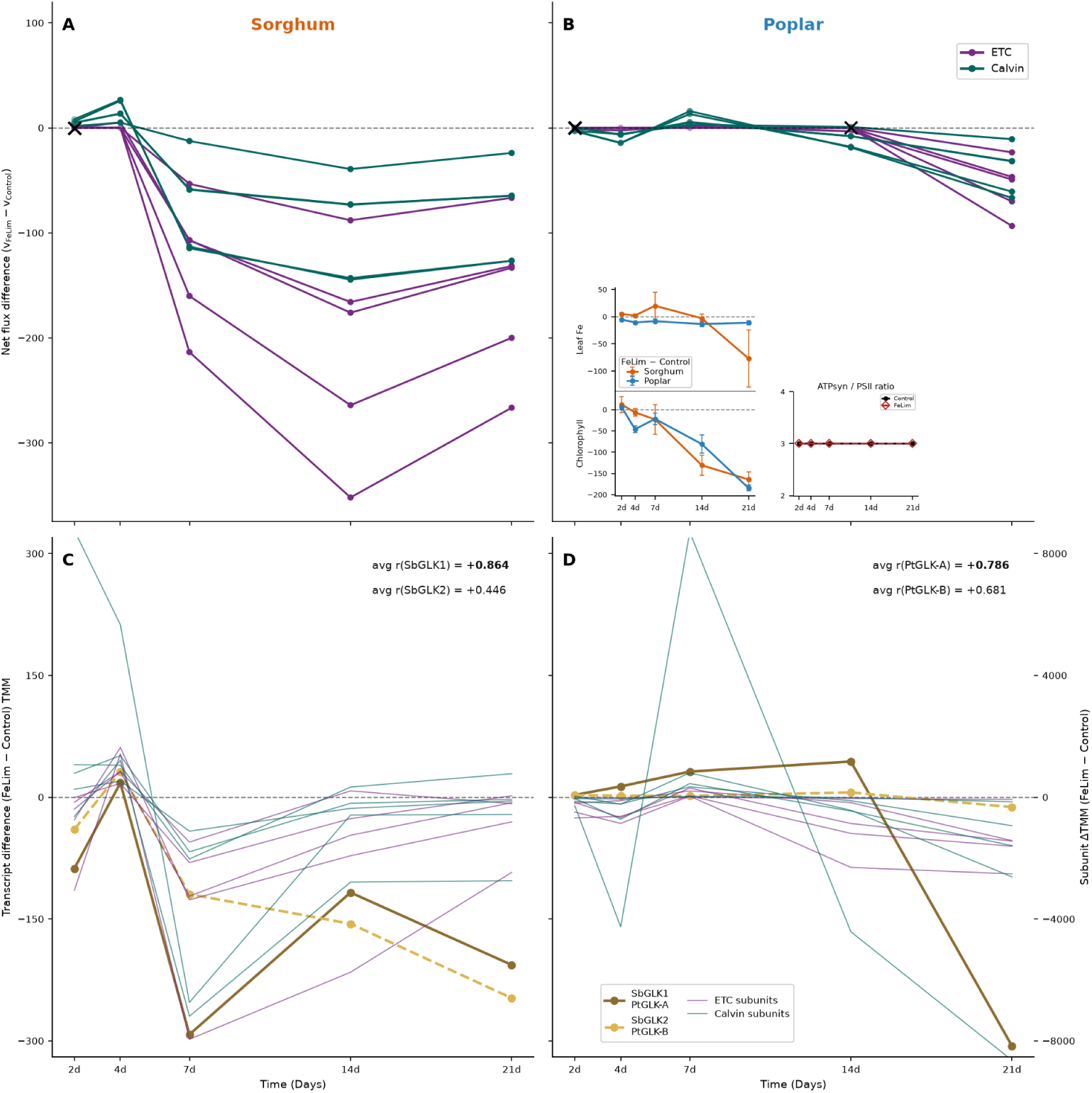
The divergent regulation of highest-flux pathways is the predominant biological signal separating the two species. A & B) Here we plot the flux through ten reactions which are the five members of the linear photosynthetic ETC — Photosystem II, cytochrome *b*_6_ *f*, Photosystem I, ferredoxin-NADP^+^ reductase (FNR), and plastidial ATP synthase — and the five Calvin-cycle enzymes RuBisCO, phosphoglycerate kinase (PGK), NADPH-glyceraldehyde-3-phosphate dehydrogenase (NADPH-GAPDH), phosphoribulokinase (PRK), and sedoheptulose-1,7-bisphosphatase (SBPase). Fluxes are reported as the FeLim-minus-Control net. The lines are unlabeled with the exception of color because the biologically meaningful signal is the mass-balanced coherence of their collective response. All ten reactions hold their direction at every timepoint under a mass-balance penalty *p* = 2 in both species. The inset on panel A shows the plastidial ATP synthase / Photosystem II flux ratio holding at a flat 3:1 across all timepoints under both Control and FeLim; it is drawn for Sorghum only, as the Poplar ratio is identical (see main text). C & D) Transcript trajectories of the two GLK paralogs per species (Sorghum SbGLK1/SbGLK2; Poplar PtGLK-A/PtGLK-B, an indistinguishable pair) compared against those of the rate-limiting subunits of each of the ten photosynthetic-regulon enzymes. The two transcription factors (TFs) are plotted on the left y-axis and the rate-limiting subunits on a twin right y-axis, because TF abundance is typically on a much smaller scale than that of their targets. The annotation in each panel gives the mean Pearson correlation of the rate-limiting subunits with each TF. Panels B and D share the y-axes of panels A and C respectively. The two insets in panel B carry the measured leaf phenotypes for both species — leaf iron content and total chlorophyll — plotted as the FeLim-minus-Control difference, so positive values mean the iron-limited leaves assayed *higher* than their controls. Poplar’s leaf iron is consistently negative throughout; Sorghum’s shows much greater replicate dispersion and no timepoint differs significantly from control. For chlorophyll Sorghum holds through day 7 and then falls steeply, Poplar declines gradually from day 4, and the two arrive at a comparable deficit by day 21.

**Figure 9.**
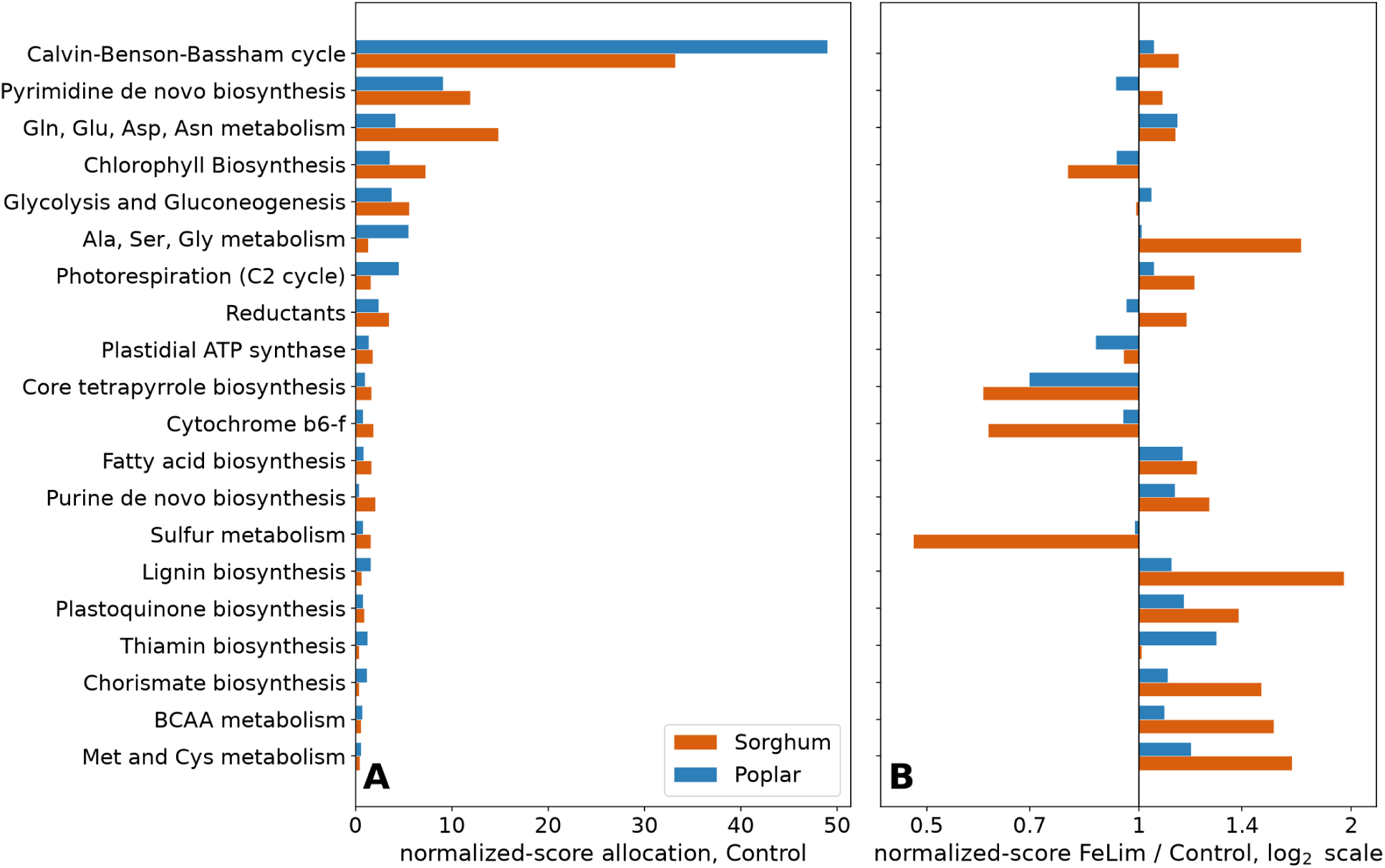
Normalization with the plastid-proteome pool reveals iron-driven reallocation. A) The share of each species’ plastid proteome devoted to each pathway under Control conditions, plotted as the percentage of that species’ total plastid transcript pool; C3 Poplar allocates a larger share to photorespiration than C4 Sorghum. B) Reallocation occurs under iron limitation; we plot the normalized-score FeLim/Control ratio, that is, the change in each pathway’s share of the pool, so that bars above 1 gain share and bars below 1 lose it. As the x-axis is logarithmic, halving and doubling are seen as equal and opposite bars. The set of subsystems is a union of the twelve subsystems with the largest Control allocation (the panel A ranking) and the twelve with the largest share change under iron limitation (the panel B ranking), restricted to subsystems holding at least 0.5% of the Control score summed over both species. As the plastid proteome contracts, core tetrapyrrole and chlorophyll biosynthesis lose share in both species (consistent with chlorosis), as do cytochrome *b*_6_ *f* and ATP synthase, while photorespiration and the Calvin cycle are protected. The eight subsystems contributed by the panel B ranking alone hold small Control allocations — short bars in panel A — but reallocate strongly; sulfur metabolism is the most extreme, with Sorghum halving its share (∼0.50) where Poplar holds its own (∼0.98). Subsystem names are abbreviated on the axis (BCAA, branched-chain amino acid; C2 cycle, photorespiration; amino acids by three-letter code).

### 3.3. Hybrid Mechanistic Solver Diagnostics

Starting from the transcript-derived vector *V_b f_* (§2.6), our gradient-descent approach converges to a set of fluxes that remains close to the nominal *V_b f_* constraints and never exceeds the post-update ceiling of 1.10*V_b f_* while substantially reducing the global mass-balance residual at every metabolite. Each experimental condition is processed independently, and the freezing mechanism (Section 2.7.2) lets the optimizer continue working on conditions that are still improving without disturbing those that have already reached a stable region.

Figure 6 characterizes optimizer behavior across the 22 Leaf-tissue conditions (11 per species: the five timepoints under each of Control and FeLim, and a shared 0hr Control baseline sampled before the treatments diverge) and across the mass-balance penalty *p*. The per-term loss trajectory shows the data and mass-balance terms descending on different timescales but reaching a stable region together; no condition’s converged flux exceeds the 1.10 ×*V_b f_* buffered clamp described in §2.7.3. The freeze-step distribution shows that conditions plateau on comparable timescales, validating a single solver run length across the experimental panel. The Pareto frontier of *L*_data_ against *L*_mass_, together with the per-condition imbalance violins and the CDF of |*SV* | per reaction, identifies *p* = 2.0 as the operating point at the elbow of the trade-off, and the triangle-split correlation heatmap shows that the converged fluxes are highly reproducible (*ρ* ≳ 0.99) across adjacent *p* values.

### 3.4. Flux Results

With the optimizer’s behavior established, we turn to the converged fluxes themselves. We first show how the biomass reaction behaves when it is not an objective (section 3.4.1), then present the dominant photosynthetic signal and its correspondence with the GLK regulon (section 3.4.2), and finally the cross-species comparison of metabolic allocation that the plastid-relative score makes possible (section 3.4.3).

#### 3.4.1. Biomass as a passive drain rather than an objective

Unlike flux balance analysis, our approach never maximizes or otherwise forces flux through the biomass reaction. The biomass reaction carries no enzyme and so is excluded when applying the data loss term; whatever flux it carries is set by mass balance alone. We do use the biomass reaction during the pFVA step to shape the feasible flux range. This affects how much flux passes through the biomass reaction rather than fixing its component coefficients, making the biomass reaction a *passive mass-balance drain*. The approach utilizes the biomass reaction as a drain, and the extent changes over the time course (Fig. 7), but only as an emergent consequence of upstream carbon supply that changes in response to limited iron.

Figure 7 tracks the carbon actually drawn into biomass over the course of the experiments. The flux through the biomass reaction is resolved by the classes of biomass components and expressed as carbon atoms (bio1 flux × carbon number). The total flux × carbon in the biomass (leftmost *Biomass* bars) is high and comparatively stable under Control in both species. Iron limitation does not depress it early— Sorghum’s FeLim biomass at 2d exceeds its own Control by a quarter, and Poplar’s tracks Control through 7d, exceeding it at 4d—but it falls steeply thereafter, and far more in Sorghum (C4) than in Poplar (C3): by 14–21d Sorghum’s FeLim biomass has collapsed to between a fifth and a third of its Control value, whereas Poplar retains roughly 70%. We cannot use the leaf iron concentrations to explain the collapse in Sorghum, because those concentrations are indistinguishable from control until the end of the time course (upper inset in Fig. 8B, and see section 3.4.2). Leaf chlorophyll does follow it: Sorghum’s falls to 0.42× its control at day 14 and 0.31× at day 21, over the same interval as the biomass collapse, whereas Poplar’s declines gradually and only reaches comparable depletion at day 21 (lower inset in Fig. 8B). This is exactly what a passive, photosynthesis-downstream drain should do: C4 carbon fixation carries higher flux and a heavier iron demand (additional Fe–S centers and electron-transport load), so restricting iron throttles Sorghum’s carbon supply, and hence its biomass drain, more sharply than Poplar’s. The biomass stoichiometry is fixed and every component class rises and falls together; sugars and amino acids are the largest carbon draws, followed by fatty acids and cell-wall components, with organic acids and nucleotides minor, and these proportions are invariant across time and treatment—only the overall magnitude changes. The whiskers on each bar mark how much carbon the rest of the network actually supplies to that class, so the gap between whisker and bar is the mass-balance slack the solver absorbs: amino acids are consistently under-supplied, meaning plastidial biosynthesis lags the demand written into the biomass reaction, while sugars and organic acids are consistently over-supplied.

The carbon is nonetheless delivered by genuine biosynthesis: every biomass component is produced by its canonical plastidial pathway—glutamate and glutamine by ferredoxin/NADH-dependent glutamate synthase and glutamine synthetase, the aspartate-family amino acids via aspartokinase and threonine synthase, aromatic amino acids via arogenate dehydrogenase, fatty acids via the acyl-ACP thioesterases, sugar phosphates via fructose-1,6-bisphosphatase, and nucleotides via the IMP and OMP pathways. Each pathway carries flux at a transcript-set rate, so all are simultaneously active and differ only in magnitude across classes, timepoints, and iron treatment.

We quantified how much of the network flux remains coupled to the transcriptome after the projection. For each reaction we correlated its converged net-flux trajectory across the ten treated Leaf conditions with its transcript-derived *V_b f_* trajectory (Section 2.6), calling a reaction *transcript-coupled* when the two agree at Pearson *r* ≥ 2/3. Weighting each reaction by its carbon throughput — peak absolute flux multiplied by the carbon atoms on its substrate side — 94% (Sorghum) and 93% (Poplar) of the carbon flux through reactions carrying a transcript target remains coupled to it, the balance being set by mass balance alone; by simple count, however, only 49% and 52% of those reactions are coupled, so the agreement is concentrated in the largest fluxes rather than spread evenly across the network. The carbon-weighted value is insensitive both to the threshold, which can be raised to *r* = 0.77 before it moves, and to the mass-balance penalty, which shifts every class by under 1.5 percentage points between *p* = 1.0 and *p* = 2.0. Because *V_b f_* is both the starting value and the ceiling, this measures how much of the transcript pattern survives being made mass-balanced rather than showing independently that transcripts set flux.

#### 3.4.2. The dominant signal: photosynthetic electron transport and Calvin cycle

The predominant biological signal separating the two species is a coordinated, species-specific perturbation of the photosynthetic apparatus and the Calvin cycle under iron limitation (Figure 8). We follow ten reactions together: the five complexes of the linear electron-transport chain (PSII, cytochrome *b*_6_ *f*, PSI, FNR, and the coupled plastidial ATP synthase) and five Calvin-cycle enzymes (RuBisCO, PGK, NADPH-GAPDH, PRK, and SBPase). These ten were chosen not merely because they carry high flux, but because they are recognized targets of the GOLDEN2-LIKE (GLK) transcription factors that coordinate the photosynthesis-associated nuclear-gene regulon (Waters et al., 2008, 2009); tracking them together therefore tests whether a single transcriptional module behaves coherently in the input data and is carried through the projection as a coordinated, mass-balanced block of flux. To ensure these responses are not artifacts of the mass-balance penalty *p*, we ran the gradient descent at four penalties (*p* ∈ {0.1, 0.5, 1.0, 2.0}) and consider a reaction’s iron response at a timepoint *defensible* when all four penalties agree on its direction, the change exceeds ∼2% of the reaction’s flux scale (roughly twice the empirical noise floor), and the local mass-balance residual stays ≤ 0.30. Nearly every datapoint across these ten reactions is defensible for both species and across the time course, so the collective signal is biologically driven even where individual per-timepoint values approach the noise floor.

Plotting the differences in flux between FeLim and control conditions across the leaf time course (Fig. 8A,B), the ten reactions move as a mass-balanced ensemble rather than independently — the behavior expected of a fully coupled subnetwork in the sense of flux coupling analysis (Burgard et al., 2004; Marashi et al., 2012). One way to verify that the solver honors this coupling is the ratio of ATP synthase to PSII. In our reconstruction, lumenal H^+^ is produced only by PSII (4 H^+^ per O_2_) and cytochrome *b*_6_ *f* (4 H^+^ per turnover) and consumed only by the plastidial ATP synthase (4 H^+^ per ATP), while plastoquinone A is exchanged only between PSII and cytochrome *b*_6_ *f* (2:1). Mass balance therefore *forces v*_ATPsyn_ = 3 *v*_PSII_. We stress that the flat 3:1 ratio we observe (inset in panel A) is consequently not an independent biological finding; it is what any mass-balanced approach would find given the stoichiometry. What is established here is that the *soft* mass-balance penalty is tight enough at *p* = 2.0 for the balance to hold across the time course in both species, despite order-of-magnitude shifts in the fluxes. Overall, the response is drastically different between Sorghum and Poplar. Sorghum shows an abrupt, deep *suppression* at day 7: the FeLim-minus-Control difference reaches its most negative value across the entire panel (down to ∼− 210 flux units at photosystem I, the most affected reaction), after which the ensemble partially recovers through days 14 and 21. Poplar shows no comparable acute event; apart from a small positive excursion at day 7 its suppression deepens steadily, and is greatest at day 21.

The two measured leaf phenotypes of iron and chlorophyll, inset in panel B, place these flux responses in context. Both report the FeLim-minus-Control *difference*, with error bars propagated from three biological replicates per treatment arm, and both carry the two species together. For iron, in Poplar the difference is small, negative and consistent throughout (Welch’s *t*, *p* < 0.05), while in Sorghum the replicate scatter is up to an order of magnitude larger and no timepoint differs significantly from control (Welch’s *t*, *p* ≥ 0.15), so we do not interpret that trajectory. Note that ICP-MS reports iron per unit dry weight, and iron-limited seedlings accumulate less biomass, so a positive difference can reflect reduced growth rather than iron sufficiency. In chlorophyll, the trend is similar between the two species: Sorghum holds through day 7 and then falls steeply, Poplar declines gradually from day 4, and the two arrive at a comparable deficit by day 21. In Sorghum the simulated flux response at day 7 therefore precedes the visible pigment loss; in Poplar the two decline together.

Overlaying the GLK transcript trajectories on those of the rate-limiting subunit of each reaction (Fig. 8C,D) reveals a species-specific pattern. This comparison is deliberately between transcripts and not simulated flux. In both species one paralog correlates more strongly with the subunits than the other, and does so for every one of the ten subunits without exception; what differs between the species is how large that separation is. In C4 Sorghum the subunits favor SbGLK1 over SbGLK2 by an average Pearson *r* of +0.86 against +0.45, which is to be expected of the sub-functionalized, bundle-sheath/mesophyll-compartmentalized GLK pair of C4 grasses (Wang et al., 2012). In C3 Poplar, the two GLK copies (PtGLK-A and PtGLK-B) are a recent duplication that cannot be orthologously separated into GLK1 and GLK2; as such, they do not exhibit the same degree of differential regulation (+0.79 against +0.68, a fourfold smaller gap).

#### 3.4.3. Relative reaction scores enable cross-species allocation

The transcript-driven flux above uses the objective reaction score *r_s_*, which is appropriate *within* a species but not *across* species, because the two plastid proteomes differ substantially in size (Table 1). Despite Poplar carrying more matched plastid orthologs than Sorghum (2,489 vs. 2,008), Sorghum’s plastid genes are expressed at roughly twice the level, so objective scores carry a species-specific scale offset (the plastid proteome pools differ ∼1.7×) and cannot be read as comparable quantities. Expressing each score relative to the summed plastid proteome *P_total_*—the plastid-relative reaction score *r̃_s_* (section 2.4)—converts abundances into a *share* of the plastid proteome, which is comparable across species and lets us interpret metabolic *allocation* rather than raw flux. As we are using identical plastidial networks for both species, we are able to compare them directly.

Two consequences follow. First, the proteome-share view exposes cross-species differences in *allocation* that the objective score cannot: C3 Poplar devotes a larger share of its plastid proteome to photorespiration than C4 Sorghum, while Sorghum allocates comparatively more to chlorophyll biosynthesis and to glutamine/glutamate/aspartate metabolism (Figure 9A). Second, because the pool contracts under iron limitation, the relative score reveals which pathways the plastid *prioritizes* as it downsizes: core tetrapyrrole and chlorophyll biosynthesis lose share in both species—consistent with the onset of chlorosis—as do cytochrome *b*_6_ *f* and ATP synthase, while photorespiration and the Calvin cycle are protected (Figure 9B).

Sulfur metabolism falls to half its Control share in Sorghum (0.50) while Poplar holds steady (0.98), and methionine and cysteine metabolism moves the other way, gaining most in Sorghum (1.63 against 1.19). Sulfur and iron metabolism are known to be interdependent in plants (Astolfi et al., 2021), and the four plastid sulfur reactions do not move together. Of 333 scored Sorghum reactions the two reductive steps rank in the bottom 2% — adenylyl-sulfate reductase (APR, EC 1.8.4.9; 0.30) and ferredoxin–sulfite reductase (SiR, EC 1.8.7.1; 0.55) — while the competing branch to PAPS, adenylyl-sulfate kinase (EC 2.7.1.25; 2.06), is among the strongest gainers. In Poplar the split is muted: SiR falls (0.73) but APR does not (1.01). Cysteine synthase (EC 2.5.1.47; Sorghum 1.96) carries most of the methionine and cysteine gain.

## 4. DISCUSSION

In this work we extend a hybrid ML–FBA framework so that it integrates transcriptomic data to constrain the internal fluxes of a metabolic network, rather than optimizing growth from media and growth-rate measurements. Flux balance analysis excels for microbial systems held in a fixed medium at steady state, but its assumptions are difficult to satisfy in the cells and tissues of complex multicellular organisms, which occupy transient, spatially heterogeneous metabolic states. We therefore turned the microbial input/output formulation inside out: instead of fixing the medium and reading out growth, we use the transcriptome — an easily sampled proxy for enzyme availability — to bound the biologically feasible flux of each reaction, and let gradient descent find a flux distribution that respects those bounds while softly constraining the stoichiometric mass balance. We tested this on the plastidial metabolism of the C4 monocot *Sorghum bicolor* and the C3 dicot *Populus trichocarpa*, across a leaf time course under iron limitation.

The defining property of the approach is that the biomass reaction does not dictate flux; the transcriptome does. The biomass behaves as a passive mass-balance drain (Figure 7) rather than an objective that pulls flux through the network. The transcripts determine where carbon flows, so the model reports variable activity in pathways *not* tied to biomass biosynthesis, and in turn allows the production of biomass and of its respective components to vary rather than being fixed by an objective. This is the central departure from any FBA approach: we allow flexible constraints to capture the transient metabolic state implied by the transcriptome. It is worth separating which layer of the framework produced each of the results that follow. The carbon drawn into the biomass drain (Figure 7) and the coordinated suppression of the electron-transport chain and Calvin cycle (Figure 8A,B) are flux-level results: how much carbon the network can actually deliver, and how the high-flux subnetwork moves as a coupled block, depend on the stoichiometry as a whole and cannot be read off any single reaction’s transcript-derived capacity. The coherence of the GLK regulon (Figure 8C,D) and the split within sulfate assimilation (Figure 9B) are results of the transcript layer — the per-reaction capacities and their pool-normalized shares — and we present them as such.

The strongest and most coherent signal to emerge was a coordinated, species-specific perturbation of the photosynthetic electron-transport chain and Calvin cycle under iron limitation (Figure 8), recovered as a mass-balanced ensemble whose rate-limiting subunits track the GOLDEN2-LIKE (GLK) regulon. The two species diverge sharply. Sorghum shows an acute, synchronized suppression that is deepest at day 7, in which the regulon follows one GLK paralog far more strongly than the other, the differentiated signature expected of the sub-functionalized, bundle-sheath/mesophyll-compartmentalized GLK pair of C4 grasses (Wang et al., 2012). Poplar responds more gradually, and its two GLK copies — an indistinguishable recent Salicaceae whole-genome-duplication pair — correlate with the regulon similarly, the redundant pattern characteristic of C3 species. That the GLK regulon and the transcripts of its photosynthetic targets move coherently, and that their species-specific pattern is consistent with the C3/C4 distinction, indicates that the transcriptomic signal driving the model carries genuine regulatory structure.

Expressing each reaction score relative to the total plastid proteome adds a perspective the objective score cannot provide (Figure 9). The two species’ plastid proteomes differ substantially in size and only the proteome-relative score makes their pathway investment comparable across species, exposing differences in metabolic *allocation*; and because the pool itself contracts under iron limitation, the relative score reveals which pathways the plastid prioritizes as it downsizes. Within Sorghum’s plastidial sulfate-assimilation pathway the share change separates by enzyme cofactor rather than by position in the pathway. The two steps that lose allocation are catalyzed by enzymes containing iron–sulfur clusters, whereas the enzyme catalyzing the step that gains allocation requires no iron. The pattern is to be expected under limited iron and places sulfate reduction along with tetrapyrrole synthesis and cytochrome *b*_6_ *f*, consistent with the intracellular ordering of iron allocation reported for iron-limited cells (Naumann et al., 2007). We stress that sulfate assimilation in FBA is pinned to the biomass demand for cysteine and methionine and cannot halve unless biomass does, and the split at the APS branch point would be left underdetermined by mass balance.

Our approach builds directly on the AMN family of neural-mechanistic hybrid models (Faure et al., 2023). Our contribution is to integrate transcriptomics-derived flux bounds directly into the gradient descent, so that the transcript signal shapes the flux distribution during optimization. The two components of our approach are well established: gating or scaling reactions by transcript abundance is a long-standing idea, from E-Flux (Colĳn et al., 2009) to transcriptome-guided parsimonious methods such as RIPTiDe (Jenior et al., 2020), as is relaxing a fixed biomass recipe through flexible biomass composition (Shaw & Kundu, 2015; Yuan et al., 2016). The novelty lies in their combination. This also sets our method apart from those expression-integration methods, which retain a growth objective and a hard steady state; and from hybrid pipelines such as FlowGAT (Hasibi et al., 2024), which first solve FBA under a growth objective and then train a neural network on the resulting flux graph. With FlowGAT an objective-driven solution precedes and constrains the learning step. The same ordering applies to machine-learning models trained to predict flux control from simulated flux distributions (Kugler & Stensjö, 2024): the mechanistic solve is upstream of the learning, whereas here the two are the same computation.

Three limitations bound what can be concluded from this work. First, and most importantly, we have no flux-level ground truth. No ^13^C metabolic flux analysis, enzyme assay, or other independent measurement of flux was available for these tissues, so the solutions are validated only by their qualitative consistency with known iron-limitation physiology and by internal checks that the solver honors stoichiometry. Second, the method inherits the assumption — shared with every transcript-integration method — that transcript abundance tracks enzyme abundance, which holds unevenly across enzymes and conditions. Third, because the transcript-derived vector is both the initial condition and the upper bound, the approach measures how the transcriptome is *reshaped* by mass balance rather than testing the transcriptome against an independent signal. Within those bounds, our method demonstrates value in studying metabolic responses in tissues of complex organisms, by accommodating the non-steady, transient states tissues can occupy, at the cost of tolerating a controlled amount of slack in the mass balance. This study is limited to plastidial metabolism, chosen for its central role in synthesizing primary biomass components and its rough comparability to a microbe, but several extensions follow naturally. First, additional omics layers — fluxomic and metabolomic data — can be incorporated as further constraints as they become available. Second, because the approach is implemented as a differentiable gradient-descent procedure in TensorFlow, it retains the capability that defines the AMN family: although the underlying QP could be solved directly, casting it as a differentiable objective lets the same mechanistic loss serve as the trainable layer of a neural network that predicts fluxes directly from omics datasets, and be extended with additional loss terms. Third, the pipeline itself can scale from plastidial metabolism to full genome-scale reconstructions.

## 5. DATA AND CODE AVAILABILITY

All source code is available in two public repositories: transcript processing and the calculation of reaction scores at https://github.com/samseaver/RNASeq_Enzyme_Abundance, and the mechanistic loss function and gradient-descent implementation at https://github.com/samseaver/Biochem_Informed_Mechanistic_Gradient_Opt. The plastidial metabolic reconstructions for both species are included, as both derive from published genomes, but the code for the annotation of the enzymes in the genomes and for the reconstruction of metabolism is found in release v2.5 of PlantSEED: https://github.com/ModelSEED/PlantSEED/releases/tag/v2.5. For Sorghum, we additionally provide the transcript-derived reaction scores and *V_b f_* vectors for all 11 leaf conditions, together with the converged flux vectors at each mass-balance penalty. The equivalent Poplar files are available from the corresponding authors on request.

